# High-throughput functional characterization of combinations of transcriptional activators and repressors

**DOI:** 10.1101/2022.12.20.521091

**Authors:** Adi X. Mukund, Josh Tycko, Sage J. Allen, Stephanie A. Robinson, Cecelia Andrews, Connor H. Ludwig, Kaitlyn Spees, Michael C. Bassik, Lacramioara Bintu

## Abstract

Despite growing knowledge of the functions of individual human transcriptional effector domains, much less is understood about how multiple effector domains within the same protein combine to regulate gene expression. Here, we measure transcriptional activity for 8,400 effector domain combinations by recruiting them to reporter genes in human cells. In our assay, weak and moderate activation domains synergize to drive strong gene expression, while combining strong activators often results in weaker activation. In contrast, repressors combine linearly and produce full gene silencing, and repressor domains often overpower activation domains. We use this information to build a synthetic transcription factor whose function can be tuned between repression and activation independent of recruitment to target genes by using a small molecule drug. Altogether, we outline the basic principles of how effector domains combine to regulate gene expression and demonstrate their value in building precise and flexible synthetic biology tools.

## Introduction

Transcription factors (TFs) and chromatin regulators (CRs) contain short effector domains that can act as repressors or activators when recruited at a target gene^1,23–51,2^. Site-specific recruitment assays of effector domains and full length TFs at reporter genes have long been used to understand their effects on gene expression and develop better tools for gene regulation ^6–15^.

Although recruitment assays have historically focused on recruiting only one transcriptional effector per cell, combinatorial function is a key property of both chromatin-mediated gene regulation^16^ and transcription factor-mediated gene regulation^17–19^. Transcription factors frequently contain multiple effector domains with potentially opposing functions, with studies reporting up to 40% of transcription factors having at least 2 distinct effector domains^2,20^. For example, the chromatin regulator MGA was recently shown to feature two repressive domains with different rates of silencing and amounts of memory^21^, while an earlier study showed that the transcription factor NIZP1 features an activating KRAB domain that is dominated by a repressive C2HR domain^22^. Combinations of effector domains have a long history in the context of synthetic biology, where the well-known transcriptional activators VP64^23^ and VPR^24^ were built via combining multiple known activation domains. Similarly, repressor combinations featuring the KRAB domain of ZNF10 and DNA methyltransferases have been shown to produce a robust combination of rapid gene silencing and long-term epigenetic memory^9,25^.

A systematic understanding of how combinations of transcriptional effector domains function in human cells would expand the range of compact tools available for epigenetic perturbations and therapy^26,27^. Additionally, composing effector domains can serve as a useful strategy to design synthetic TFs capable of implementing gene regulatory functions not achievable by fusing individual effectors to DNA-binding domains. These TFs could be used for many applications, including more efficient reprogramming of cell lineage specification or high-throughput screening of the noncoding genome^28,29^.

In order to systematically test combinations of effector domains, we need high-throughput methods: even testing all possible combinations resulting from pairing 100 effector domains requires 10,000 measurements. Recently, pooled screens have been developed for high-throughput characterization of individual transcription factors and effector domains in yeast^30,31^, drosophila^32^, and human cells^20,21,33,34^. Arrayed high-throughput measurements of combinations of chromatin regulators in concert with VP16 have been performed in yeast to test 223 combinations^7,14^ and low-throughput measurements have been performed in human cells^35^. However, we are not aware of any systematic high-throughput studies to date in mammalian cells that measure how effector domains act in combination to regulate gene expression. As such, it remains unclear whether pairs of activator domains synergize when combined, whether pairs of repressor domains do the same, and how activator and repressor domains affect each other when recruited simultaneously. Here, we modify a recently developed pooled high-throughput method^21^ to test thousands of combinations of previously characterized protein domains that can activate and repress gene expression in human cells in order to start addressing these questions.

### Developing a workflow for combinatorial screening of transcriptional effector domains

We began by selecting a panel of effector domains and controls from our previous screen of single domains^21^: 44 repressors, 30 activators, and 20 control domains (**Fig. 1A**, **Materials & Methods, Supplementary Table 1**). These effectors were chosen to span a wide range of individual activation and repression strengths when recruited at the same reporter gene, (**Fig. S1A**). The control domains were chosen to be either random sequences or fragments of the DMD protein that were shown to have no effect on gene expression when recruited to a reporter individually^21^. To avoid problems with protein stability, we chose only well-expressed, stable domains as measured by FLAG-staining^21^. We generated combinations of these individual domains using a 2-step cloning strategy to build a pool of domain-linker-domain concatenations that were then cloned into a lentiviral backbone vector as fusions to the reverse TetR (rTetR) inducible DNA binding domain (**Fig. 1A**, **Fig. S1B, Materials & Methods**). We delivered this pool into K562 reporter cells using lentivirus with a low multiplicity of infection (MOI=0.24), such that most cells expressed a single library element. We measured the ability of each pair in this library to activate or repress a reporter gene using a high-throughput pooled method we recently developed: HT-recruit^21^. Briefly, by adding doxycycline (dox) we recruited the rTetR-concatenated fusions to either a minimal promoter to measure activation, or to a highly expressed constitutive promoter to measure repression (**Fig. 1B**). We separated cells into populations with high (ON) and low (OFF) reporter gene expression using magnetic separation (**Materials & Methods**, **Fig. 1C**, **Fig. S1C**), and computed the relative enrichment of individual effector combinations in each population (**Fig. 1D**). We recovered the large majority of concatenations that were present in our combinatorial library with sufficient sequencing depth (**Fig. S1D**), and found good correlation across replicates for both our activation (**Fig. 1E**, Pearson R=0.803, p<2.2×10^-16^) and repression (**Fig. 1F**, Pearson ρ=0.795, p<2.2×10^-16^) screens. To identify effector combinations for which we could reliably measure activity, we used the distribution of negative control-only combinations: we considered a pair to be activating and/or repressing if it had a score at least 2 standard deviations away from the mean of the negative control combinations.

**Figure 1.**
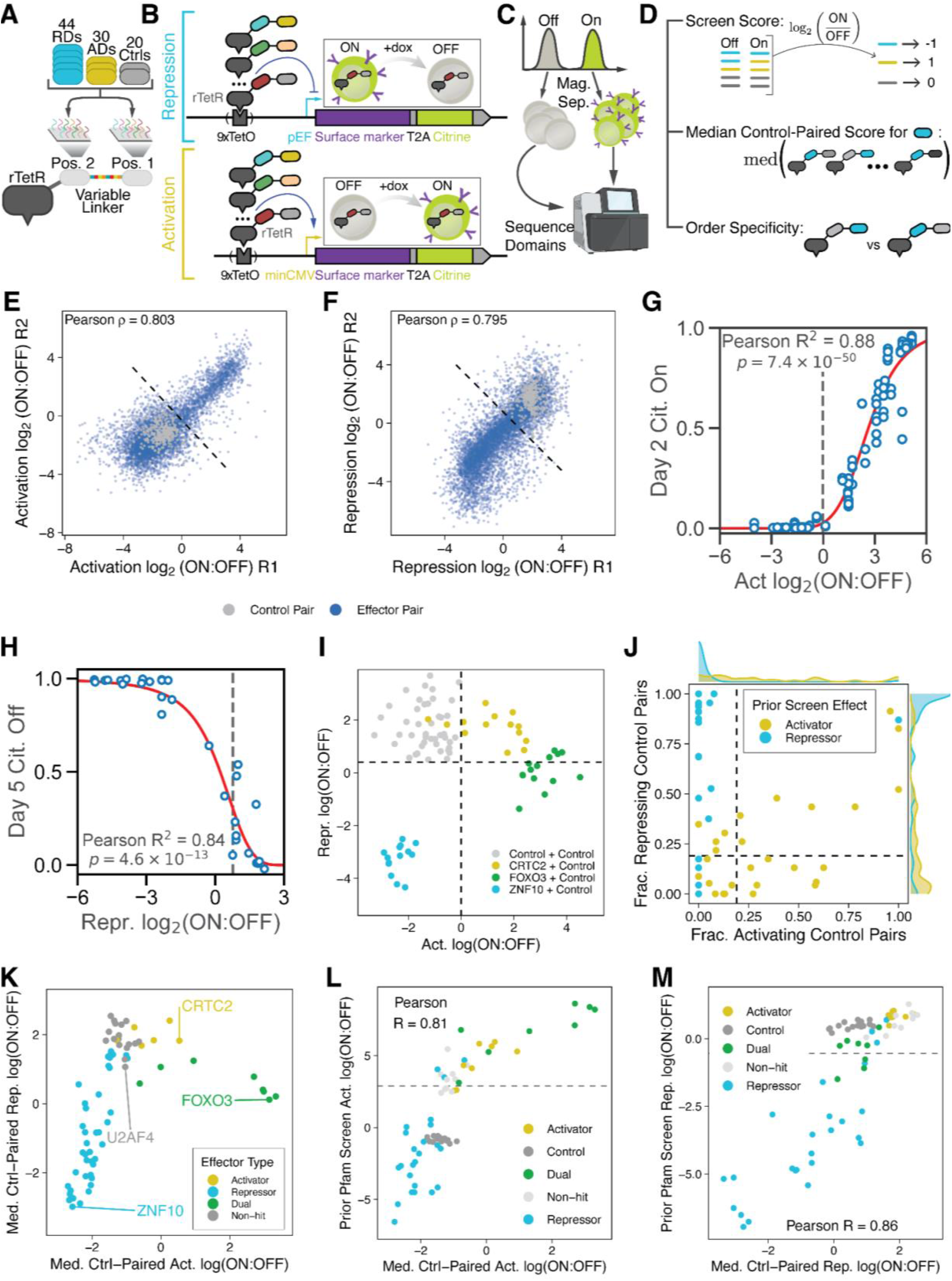
Combinatorial screening of transcriptional effector domains. A. Construction of the combinatorial library. A library consisting of approximately 100 effector domains was cloned into a lentiviral backbone in two positions (Pos. 1 and Pos. 2), connected by a 16-amino acid XTEN linker featuring a varying DNA but constant amino acid sequence, and fused to the DNA binding domain rTetR (**Materials & Methods**). This library consisted of domains that were previously identified as repressive domains (RDs), activator domains (ADs) and controls with no effect on transcription (Ctrls). B. Schematic of synthetic reporter system. rTetR-fused domain pairs are recruited to a synthetic reporter expressing a surface marker exposing an IgG epitope and an mCitrine fluorophore. The reporter gene’s promoter can be varied, with pEF being used to measure repression (top, blue arrow) and minCMV for activation (bottom, yellow arrow).. C. Magnetic separation. Cells are bound with magnetic beads that recognize the reporter surface marker and are used to separate ON and OFF populations. The DNA encoding for domains expressed in cells from each population are then sequenced via next-generation sequencing. D. Computation of screen measurements. Domain pairs are assigned a screen score equal to log_2_(ON:OFF) where ON is the relative proportion of a domain in the ON population and OFF is the relative proportion of a domain in the OFF population. The median control-paired score for a given domain is computed as the median score of all pairs featuring that domain and a negative control domain. Order specificity was determined by determining if the domain function significantly differently when positioned upstream or downstream of the negative control domain. E. Correlation between 2 replicates for the activation screen measurements for all domain pairs after 2 days of recruitment at the minCMV promoter, with domains colored based on whether they are composed of 2 negative control domains (gray) or at least 1 effector domain (blue). Dashed line is the threshold for a combination being labeled as activating, 2 standard deviations away from the mean of the negative control-negative control combinations. F. Correlation between 2 replicates for the repression screen measurements for all domain pairs after 5 days of recruitment at the pEF promoter, with domains colored based on whether they are composed of 2 negative control domains (gray) or at least 1 effector domain (blue). Dashed line is the threshold for a combination being labeled as repressing, 2 standard deviations away from the mean of the negative control-negative control combinations. G. Correlation between activation log_2_(ON:OFF) scores (x-axis) and the fraction of cells activated after 2 days of doxycycline recruitment to the weak minCMV promoter (y-axis) during individual low-throughput measurements of each pair’s function via flow cytometry. The sigmoid curve represents the best-fit line of the form y = 1/(1 + (x/k)^n). Vertical dashed line corresponds to the activation threshold for effector domain combinations as defined above. H. Correlation between repression log_2_(ON:OFF) scores (x-axis) and the fraction of cells repressed after 5 days of doxycycline recruitment to the strong pEF promoter (y-axis) during individual low-throughput measurements of each pairs’s function via flow cytometry. The sigmoid curve represents the best-fit line of the form y = 1 - 1/(1 + (x/k)^n). Vertical dashed line corresponds to the repression threshold for effector domain combinations as defined above. I. Variation in control-paired measurements. The log_2_(ON:OFF) scores for activation (x-axis) and repression (y-axis) is shown for negative control-negative control domain pairs in any orientation (gray), CRTC2-negative control domain pairs in any orientation (yellow), FOXO3-negative control domain pairs in any orientation (green), and ZNF10 KRAB-negative control domain pairs in any orientation (blue). Dashed lines represent activation and repression thresholds as defined above. J. Number of control-including concatenations that exceed the threshold for activation (x-axis) and repression (y-axis) for all domains, colored by whether that domain acts as a repressor (blue) or activator (yellow) when recruited alone. Horizontal and vertical dashed lines indicate thresholds needed for a domain to be called as an activator or repressor. K. Median control-paired activation (x-axis) and repression (y-axis) scores for individual effector domains. Activators are colored yellow, repressors blue, dual-functional effectors green, and non-hits gray. L. Correlation between activation scores from the current domain combination screen and prior nuclear Pfam single-domain activation screen. Domains present in this screen that were measured for activation in a prior screen are shown, with the median control-paired score in this screen shown on the x-axis and the log_2_(ON:OFF) score from the prior activation screen shown on the y-axis. The type of effector (activator, repressor, dual, negative control, non-hit) is indicated by the color of each point. Dashed line indicates the threshold for activation in the prior activation screen. M. Correlation between repression scores from the current domain combination screen and prior nuclear Pfam single-domain repression screen. Domains present in this screen that were measured for repression in a prior screen are shown, with the median control-paired score in this screen shown on the x-axis and the log_2_(ON:OFF) score from the prior activation screen shown on the y-axis. The type of effector (activator, repressor, dual, negative control, non-hit) is indicated by the color of each point. Dashed line indicates the threshold for repression in the prior activation screen.

To validate these high-throughput measurements and our detection threshold, we measured activation for 61 domain concatenations and repression for 38 domain concatenations by recruiting them individually to our synthetic reporters and measuring gene expression via flow cytometry. We measured the fraction of cells activated at the weak minCMV promoter after 2 days of recruitment and found strong correlations between our screen measurements and the fraction of cells activated (**Fig. 1G**, R^2^=0.88, p=7.4×10^-50^). We found a similarly strong correlation between repressive screen scores and the fraction of cells silenced after 5 days of recruitment to the strong pEF promoter (**Fig. 1H**, R^2^=0.84, p=4.6×10^-13^). We concluded that our screen accurately measured these concatenations’ ability to either activate or repress gene expression.

We began our analysis of the screen data by examining the behavior of effector domains in our library in combination with the negative control domains. We identified 4 out of 20 negative control domains that significantly affected either activation or repression scores across all effector domains and filtered them from downstream analyses (**Fig. S1E-F**). We then sought to determine if the order of the domains within concatenations altered their effect on gene expression. While prior literature has demonstrated that the ordering of effector domains in synthetic TFs can significantly affect the function of fusions^24,25^, we found that few of the domains in our library changed effect size significantly when switched from the first to the second position across all of their fusions with negative controls (**Fig. S1G**). We excluded concatenations featuring any domain in the orientation that ablated its function from downstream analysis (**Fig. S1H**).

To check to what extent effectors maintained their function when fused to a control sequence, we examined the distribution of scores for each effector domain when paired with all negative controls (**Fig. 1I**). While for some domains all scores clustered together in one quadrant (e.g., the strong KRAB repressor domain from ZNF10, **Fig. 1I**, blue), we found that other domains dropped under the detection threshold when paired with certain negative controls (e.g., weak CRTC2 activation domain, **Fig. 1I**, yellow). Some domains, such as the FOXO3 activation domain, have been previously reported to act as dual effectors that both activate a minimal reporter and repress a constitutive one when recruited individually^20^. We indeed found that most FOXO3-control fusions acted as dual activator-repressors, though a minority of fusions only acted as activators **(Fig. 1I**, green). In order to classify each domain, we calculated the number of effector-control pairs that met our hit threshold for activation and for repression (**Fig. 1J**). We used these data, along with the magnitude of activation and repression scores for each effector when paired with negative controls, to label effectors as activators, repressors, dual-functional effectors (both activators and repressors), or non-hits (**Fig. 1K**, **Materials & Methods**). We found that the median activation or repression score of each effector domain when paired with negative control domains correlated well with prior screen measurements of activation (**Fig. 1L**, Pearson R=0.81, p<2.2×10^-16^) and repression (**Fig. 1M**, Pearson R=0.85, p<2.2×10^-16^). Convinced that we could appropriately characterize how individual effectors behave when paired with negative controls, we proceeded to analyze the behavior of effector domain combinations with each other.

### Weak activator domain pairs synergistically drive gene expression from weak promoters

The high-throughput measurements for transcriptional activation identified a large number of activator-activator pairs whose activation scores exceeded the sum of each individual activator’s scores when paired with negative controls (**Fig. 2A**, left). We proceeded to validate individual examples of activator pairs to verify that the scores in the screen correlate well with individual flow cytometry measurements (**Fig. S2A-B, Fig. 1G)**, and that synergy could be replicated in low-throughput (**Fig. 2B** top). Notable examples of synergy included the pair of ANM2’s SH3 domain and KIBRA’s WW-1 domain, as well as ANM2’s SH3 domain and NOTC2’s LNR-2 domain. While the full role of the ANM2 SH3 domain is not yet clear^36^, it is known to regulate PRMT1 activity in a methylation-dependent fashion^37^, is required for the function of the actin nucleator CobI^38^, and modulates alternative splicing of BCL-X^39^. KIBRA’s WW-1 domain binds PPxY motifs in other proteins^40^, and is essential for KIBRA-mediated regulation of Hippo signaling via interactions with LATS1/2^41^, while NOTC2’s second Lin-12/Notch repeat (LNR) domain sits within its negative regulatory region^42^ and is cleaved off during ligand binding^43^.

**Figure 2.**
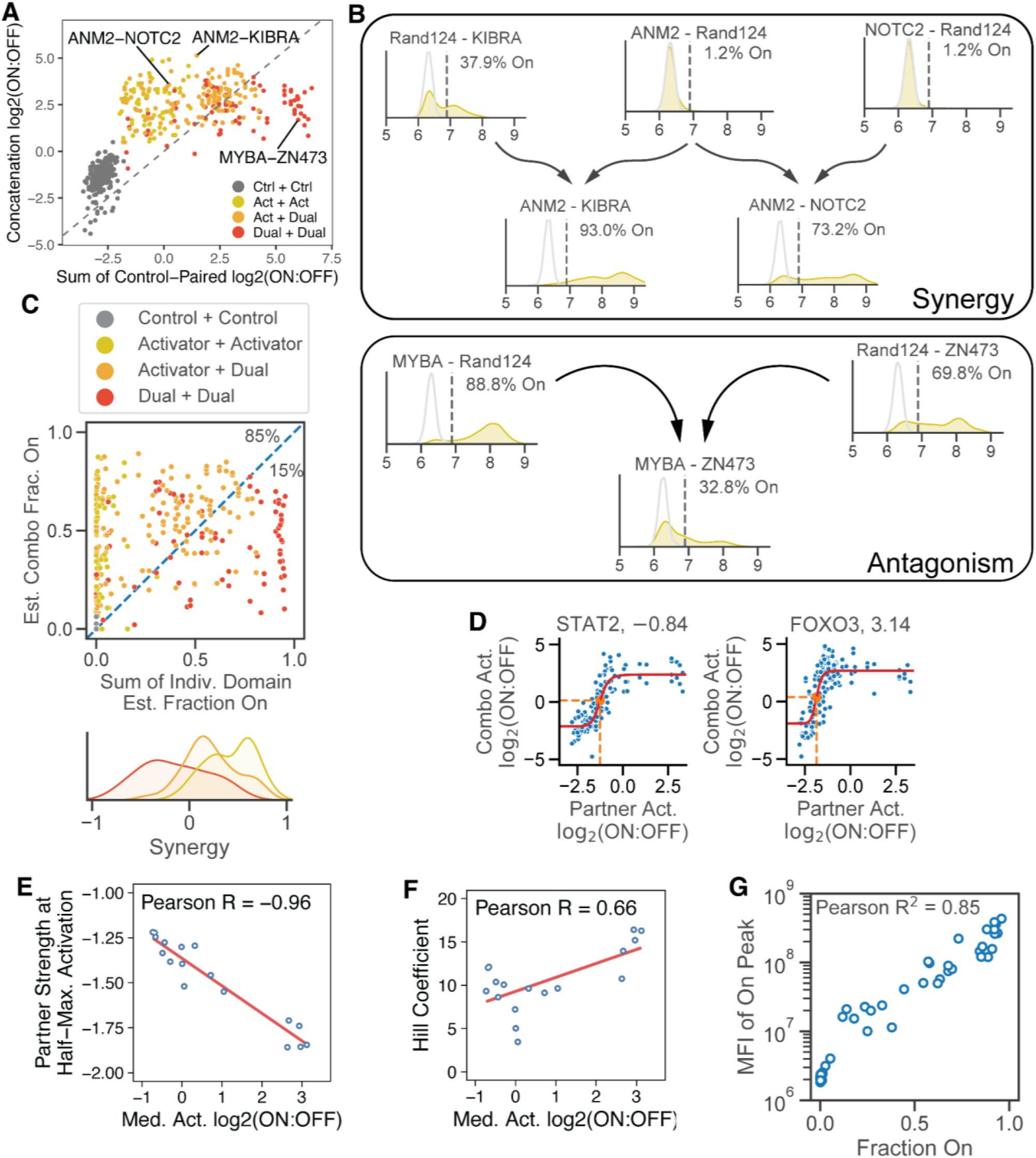
Activators work synergistically when paired together. A. Evidence for activator-activator synergy in the high-throughput screen after two days of recruitment at the minCMV minimal promoter. For negative control-negative control (dark gray), activator-activator (yellow), activator-dual (orange), and dual-dual (red) domain pairs, the sum of their individual control-paired scores (x-axis) is plotted against the average score of the combination (y-axis). Individual examples of pairs highlighted in subsequent panels are labeled and indicated by lines. B. Example low-throughput validations of activator-negative control and activator-activator pairs at the minCMV promoter. No doxycycline recruitment is shown in light gray, while the probability density of fluorescence after 2 days of dox recruitment is shown in yellow. Vertical dashed lines indicate the threshold for calling cells ON or OFF, with the fraction of cells ON in the dox population labeled within each histogram. C. Top: estimated fraction of cells activated at day 2 for every activator-activator (yellow), activator-dual (orange), and dual-dual (red) combination (y-axis) versus the sum of the component domains’ individual estimated fraction of cells activated based on control-paired scores (x-axis). Bottom: density plot of degree of synergy (x-axis) for activator-activator (yellow), activator-dual (orange), and dual-dual (red) combination pairs. Synergy is defined as the difference between the combination’s estimated day 2 fraction off and the sum of the individual domains’ day 2 fractions off. D. Sigmoidal shape of activator behavior in the high-throughput screen. For STAT2 (left) and FOXO3 (right), the control-paired score of its partner in any given pair (x-axis) is plotted against the score of the combination of the domain and that partner (y-axis) for all concatenations featuring the given domain. E. For all activator domains, the control-paired activation strength of the activator (x-axis) is plotted against the half-maximal point of the corresponding sigmoid fit to screen data (y-axis). Line shown is the line of best fit. F. For all activator domains, the control-paired activation strength of the activator (x-axis) is plotted against the Hill coefficient of the corresponding sigmoid fit to screen data (y-axis). Line shown is the line of best fit. G. Correlation between the fraction of activated cells (x-axis) and the mean fluorescence intensity (MFI) of activated cells (y-axis) for each combination measured individually.

Surprisingly, we found that the strongest activators at minCMV, which were generally dual-functional effectors that also repressed gene expression at pEF, resulted in lower activation scores at minCMV when paired together than when either activator was paired with negative controls (**Fig. 2A**, upper right quadrant). To verify that the antagonism of dual-dual effector pairs was real and not simply a result of promoter saturation, we individually tested the MYBA-ZN473 pair and found that, indeed, the combination did not activate as many cells as either domain when paired with a negative control (**Fig. 2B** bottom). Using the previously computed functions connecting screen scores with individual flow cytometry measurements of gene activation (**Fig. 1G**), we compared the estimated fraction of cells activated for each combination of activators with the sum of each activator’s control-paired estimated fraction activated. We found that activator-activator pairs tended to synergize, activator-dual pairs were additive, and dual-dual pairs acted antagonistically with low activation scores (**Fig. 2A**, **Fig. 2C** top). We computed an estimated quantity of synergy for each combination by taking the difference between that combination’s estimated fraction on and the sum of its individual domains’ fractions on. Doing so, we found that activator-activator pairs tended to feature more synergy than activator-dual pairs, which in turn tended to feature more synergy than dual-dual pairs (**Fig. 2C** bottom).

For each activator or dual-functional domain we then tracked the activity of all combinations containing that domain (including combinations with repressors and controls) and examined how the strength of the combination varied with the strength of the partner’s activation score. We fit these data using a sigmoidal Hill function (**Fig. 2D**, **Fig. S2C**). We found that the strength of the partner domain at the half-maximal point of the Hill function decreased as the activation domain’s strength increased, meaning that strong activators were able to reach half-maximal activation with less help from their partner domain than weak activators (**Fig. 2E**). We also found that the Hill coefficient of these functions went up as the activation domain’s strength increased, indicating that the increase from minimal to substantial gene activation happened more rapidly in stronger activation domains as the strength of their partners increased (**Fig. 2F**).

Across all individual validations, we found a strong correlation between the fraction of cells activated and the mean fluorescence intensity (MFI) of activated cells for our effector domain pairs (**Fig. 2G**). This correlation and the fluorescent distributions of reporter expression (**Fig. 2C**, **Fig. S2A**) are consistent with activation domains modulating transcriptional bursting kinetics at the minCMV promoter: either increasing burst frequency or burst magnitude^44–48^. Since the degradation rates of our reporter mRNA and protein are slow, taking multiple days to dilute out^10,21^, reporter molecules in our cells persist long after the cessation of a transcriptional burst. As such, it would be difficult to determine whether reporter protein molecules were made all at once in one big burst or bit by bit (in many small bursts) over a longer period of time. Thus, in this regime of slow degradation rates, increasing either the duration of a transcriptional burst (burst size) or the frequency of such bursts (burst frequency) would lead to both a higher fraction of cells measured to be on and a greater MFI of those on cells^49^. Our findings are consistent with recent work showing that changes in burst kinetics can combine to produce transcriptional synergy^50^.

### Repressor domain combinations generate robust promoter silencing

Our measurements of transcriptional repression suggested a linear relationship between the repressive strength of each domain and the repressive strength of the pair (**Fig. 3A,** points on the diagonal in the left bottom quadrant). In this case, the points that are above the diagonal come from saturation of repression at our reporter; effector pairs with scores ≤-3 are predicted to silence all cells according to our validation curve (**Fig. 1H**, **Fig. S3A**). Indeed, when we tested a number of domain pairs in low throughput, we found that recruitment of pairs of strong repressors that each fully repressed the reporter when paired with controls (e.g. ZNF10 and CBX1) can also repress 100% of the cells when the pair is recruited. (**Fig. 3B**).

**Figure 3.**
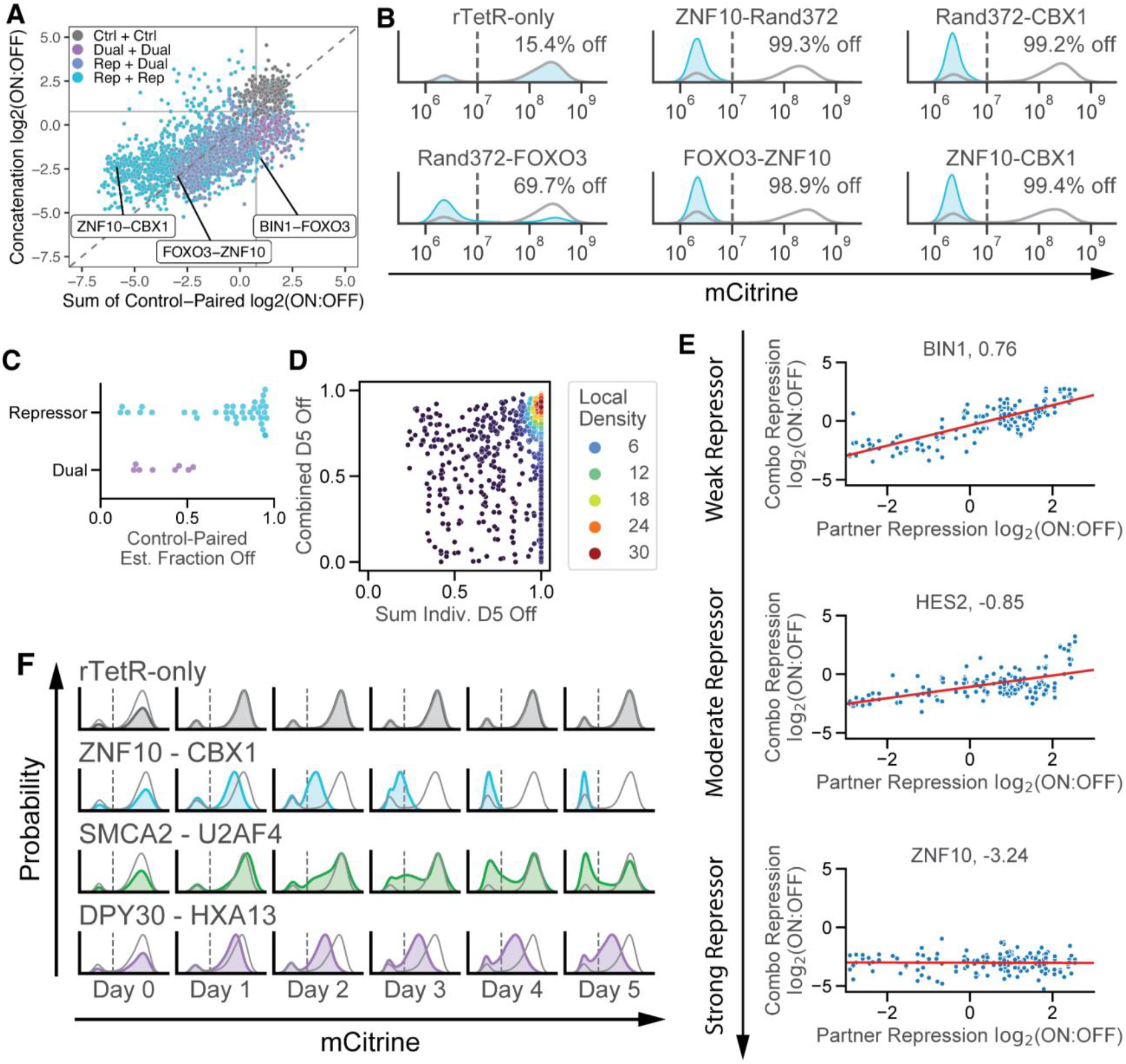
Repressors combine to drive gene silencing. A. Repressor-repressor pair behavior in the high-throughput screen after five days of recruitment at the pEF promoter. For negative control-negative control (dark gray), repressor-repressor (light blue), repressor-dual (dark blue), and dual-dual (purple) domain pairs, the sum of their individual control-paired scores (x-axis) is plotted against the average score of the combination (y-axis). Horizontal and vertical lines denote the repression threshold as defined in **Fig. 1F**, while the diagonal dotted line corresponds to the line y=x. B. Fluorescent distributions of the citrine reporter gene from low-throughput flow cytometry measurements of repressor function of various domain combinations after 5 days of recruitment at the pEF promoter, with Rand372 denoting a negative control domain. Blue: 1000 ng/mL doxycycline; gray: no dox. Vertical dashed lines indicate the threshold for calling cells ON or OFF, with the fraction of cells OFF in the dox population labeled within each histogram. C. Estimated fraction of cells silenced at day 5 (x-axis) for repressors (blue) and dual-functional domains (purple) when paired with negative controls. D. For repressor-repressor, repressor-dual, and dual-dual domain pairs, the sum of the estimated day 5 fraction off for both individual domains (x-axis) is plotted against the estimated day 5 fraction off for the pair (y-axis). Color indicates density as determined by a Gaussian kernel density estimate. E. Linear shape of repressor behavior in the high-throughput screen. For a given domain (BIN1 - left, HES2 - middle, ZNF10 - right), the control-paired score of its partner in any given pair (x-axis) is plotted against the score of the combination of the domain and that partner (y-axis) for all concatenations featuring the given domain. Red lines indicate linear fits to the data. F. Individual plots of transcriptional repression. Probability density (y-axis) plots of mCitrine-A levels (x-axis) presented for four 4 different pairs of domains (rows) as measured by flow cytometry over 5 days of doxycycline recruitment (columns).

Using the best-fit curve for low-throughput validations mapping the repression log_2_(ON:OFF) scores to the fraction of cells silenced at day 5 (**Fig. 1H**), we estimated the fraction of cells silenced at day 5 for each repressor or dual effector in our screen based on its median control-paired repression log_2_(ON:OFF) score. We found that while the strength of effectors that could repress gene expression spanned a wide range, the majority of such domains were estimated to silence ≥75% of cells after 5 days of recruitment (**Fig. 3C**). As a result, when comparing the estimated fraction of cells silenced at day 5 for a combination versus the sum of the individual domains’ respective fractions, we found that most concatenations were expected to and did silence virtually all cells after 5 days, with stronger repressors more consistently producing more repressive combinations than weaker repressors (**Fig. 3D**, **Fig. S3B-D**).

Our high-throughput measurements indicated that the repression log_2_(ON:OFF) score of a repressor domain pair was a linear function of the two individual domains, as opposed to the nonlinear and sigmoidal behavior of activation domain pairs. To determine if this feature held true for individual repressor domains, we plotted the strength of every combination featuring each domain versus the strength of the domain’s partner in that combination. We found that the trendline for repressors was linear, with lower slope for stronger repressors (**Fig. 3E, Fig. S3E**). For each repressor domain, we determined the slope and y-intercept of its corresponding trendline, and found that stronger repressors featured both a flatter slope closer to 0 (**Fig. S3F,** Pearson R=0.87, p<2.2×10^-16^) and a lower y-intercept (**Fig. S3G**, Pearson R=0.92, p<2.2×10^-16^). These results suggest that weak repressors are tunable in that the strength of a pair featuring a weak repressor can vary over a wide range depending on the strength of the partner; however, pairs featuring a strong repressor will generally only act as strong repressors.

When examining the fluorescence distributions of our reporter throughout silencing, we found that most domain combinations silenced either all or none of the cells by day 5 (**Fig. 3F**, top 2 rows), as expected from previous experiments on a small number of chromatin regulators^10^. However, a minority of combinations either silenced only a fraction of cells or reduced gene expression without fully silencing cells (**Fig. 3F**, bottom 2 rows). We attempted to see if these results were consistent with prior models of stochastic gene silencing by extending a mathematical model of gene expression we have used before to understand chromatin regulation^49^. We did so by incorporating parameters for background silenced cells at the beginning of the timecourse, basal gene expression from the pEF promoter, lag time prior to silencing, the rate of protein decay upon silencer recruitment, and the rates of gene silencing and reactivation (**Fig. S4A**, **Materials & Methods**). After fitting this model to our experimental data, we found that it was able to accurately represent the dynamics of gene silencing for both strong silencers such as ZNF10-CBX1 (**Fig. S4B**) and weaker silencers such as SMCA2-U2AF4 that only silenced the reporter in a fraction of cells (**Fig. S4C**). For these types of domains, the rates of silencing predicted by this extended telegraph model for the individual validations where the model was able to accurately fit the data, and we found a good correlation with both the screen data (**Fig. S4D**, R^2^=0.73, p=1.20×10^-9^) and the fraction of cells silenced at day 5 (**Fig. S4E**, R^2^=0.79, p=2.94×10^-11^). Silencing rates predicted by the model suggested that the rate of silencing of a combination may be linear in the sum of the silencing rates of the individual domains (**Fig. S4F**), although the correlation between screen data and silencing rates was not strong enough for a more definitive conclusion.

However, we found that our model of all-or-none gene silencing could not capture the dynamics of silencers that reduced but did not fully ablate gene expression by day 5, such as DPY30-HXA13 (**Fig. S4G**), which featured a domain from DPY30 that worked as an activator on its own in prior screens and a repressor homeodomain from HXA13^21^. For these silencers, their effect on gene expression is better explained by a decrease in the production rate in the active state, rather than transitioning to a fully silent state. Altogether, these results suggest that while strong silencers in combinations will rapidly and fully silence gene expression in most cases, weaker repressors can produce more complex dynamic patterns of gene expression.

### High-throughput profiling of activator-repressor interactions

To identify principles underlying activator-repressor interactions, we looked at all pairs featuring an activator domain in combination with a repressor domain and determined if that pair functioned as an activator only pair, repressor only pair, dual-functional pair, or non-hit pair (**Fig. 4A**, **Fig. S5A**). We found that almost all pairs featuring moderate to strong repressors tended to behave exclusively as repressors, reducing gene expression at the strong pEF promoter while failing to drive transcription at the weak minCMV promoter (blue right region, **Fig. 4A**). Domains that were dual-functional when paired with negative controls (e.g., MYBA, FOXO3, and SERTAD2) tended to be dual-functional in combinations with either weak repressors or other dual effectors (green region, **Fig. 4A**), although they were dominated by strong repressors (e.g. ZNF10).

**Figure 4.**
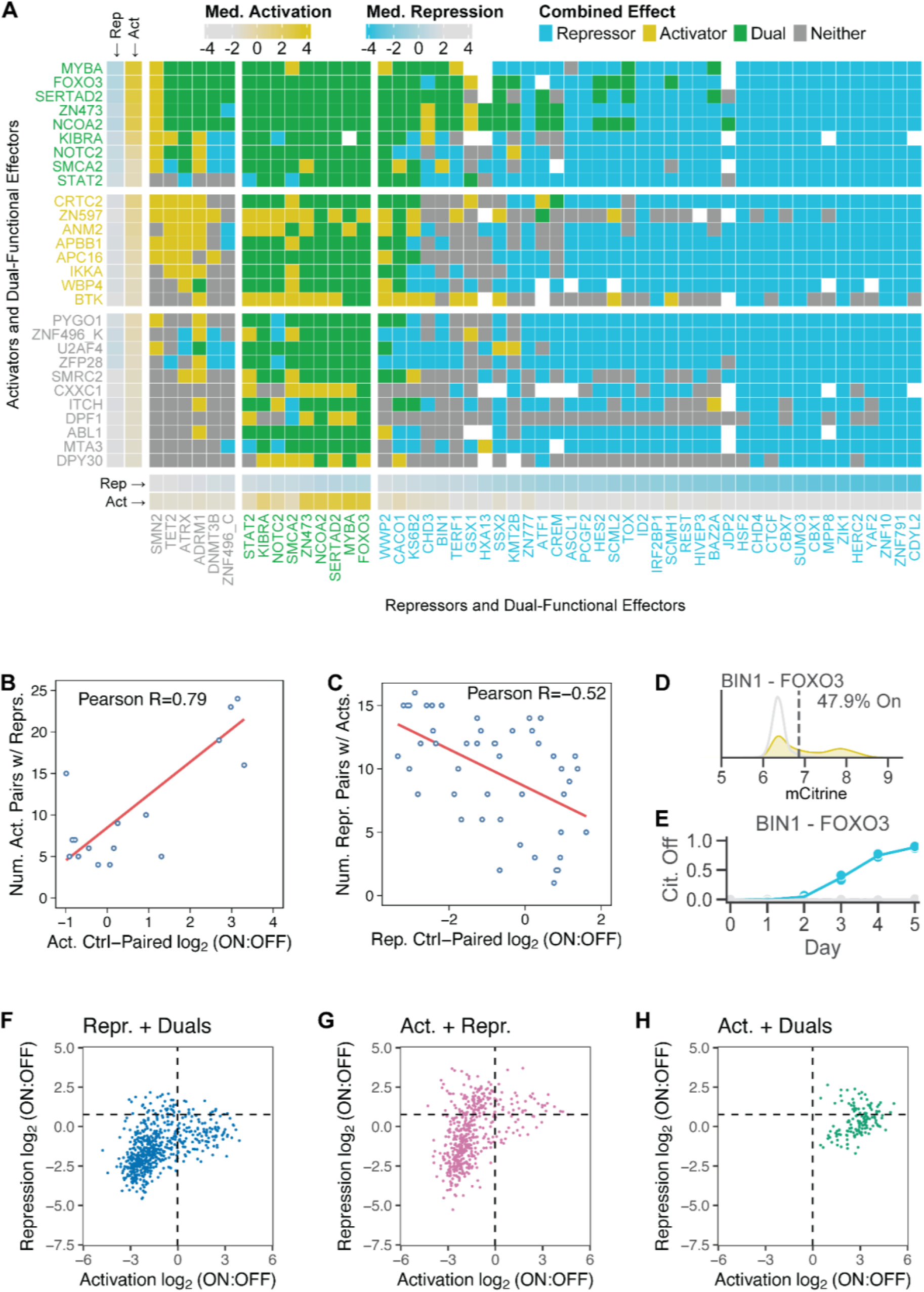
Activator-repressor interactions. A. Representation of activator-repressor interactions measured by the high-throughput screen. Domains that were identified as repressors when recruited alone are arrayed along the x-axis separated into 3 categories based on their effect when paired with controls (non-hits on the left, dual-functional domains in the middle, and repressors on the right) with the stronger repressors within each category, as measured by their control-paired repression log_2_(ON:OFF), on the right. Boxes just above each repressor domain indicate the control-paired activation/repression score for each effector. Domains that were identified as activators are arrayed along the y-axis, separated into 3 categories based on their effect when paired with controls (non-hits on the bottom, activators in the middle, and dual-functional domains on the top) with the stronger activators in each category, as measured by control-paired activation scores, on the top. Boxes just to the right of each activator domain indicate the control-paired activation/repression score for each effector. Boxes in the interior represent combinations of the corresponding activator and repressor, and are colored based on whether the combination acts as a repressor only (blue), activator only (yellow), both a repressor and an activator (green), or neither (gray). B. Correlation between the control-paired activation score for a set of activators (x-axis) and the number of pairs including any repressor domain and that activator domain that were called as an activator pair hit (y-axis). Red line represents the line of best fit. C. Correlation between the control-paired repression score for a set of repressors (x-axis) and the number of pairs including any activator domain and that activator domain that were called as a repressor pair hit (y-axis). Red line represents the line of best fit. D. Low-throughput measurement of the activation strength of the pair of BIN1’s SH3 domain and FOXO3’s TAD when recruited at the minCMV for 2 days. The gray line represents the no-doxycycline condition, and the yellow 2 days doxycycline recruitment. The vertical dashed line indicates the threshold for labeling a cell as ON or OFF. E. Fraction of cells with pEF promoter silenced by the fusion between BIN1’s SH3 domain and FOXO3’s TAD over 5 days of recruitment. The fraction of cells silenced was normalized to a no-dox condition (**Materials & Methods**). F. Distribution of repression log_2_(ON:OFF) scores (y-axis) versus activation log_2_(ON:OFF) scores (x-axis) for combinations featuring 1 repressor domain and 1 dual-functional domain. G. Distribution of repression log_2_(ON:OFF) scores (y-axis) versus activation log_2_(ON:OFF) scores (x-axis) for combinations featuring 1 repressor domain and 1 activator domain. H. Distribution of repression log_2_(ON:OFF) scores (y-axis) versus activation log_2_(ON:OFF) scores (x-axis) for combinations featuring 1 activator domain and 1 dual-functional domain.

For most domains their activator-repressor and repressor-activator orientations had similar strengths for both activation (**Fig. S5B**, Pearson R=0.82, p<2.2×10^-16^) and repression (**Fig. S5C** Pearson R=0.79, p<2.2×10^-16^), indicating that these results were not the result of position-specific effects of the protein domains. Activator-repressor combinations with stronger activators were significantly more likely to be able to activate the minimal promoter than combinations with weaker activators (**Fig. 4B**, Pearson R=0.79, p=1.6×10^-4^). For repressors, this relationship was significantly less pronounced: weak repressors were only slightly less effective than strong repressors when paired with activator domains (**Fig. 4C**, Pearson R=-0.52, p=1.6×10^-4^). Testing pairs in low throughput validated that the combination of the weak repressive SH3 domain from BIN1 and FOXO3’s TAD could both activate gene expression at the weak minCMV promoter (**Fig. 4D**) and repress gene expression when recruited to the strong pEF promoter (**Fig. 4E**). As expected, adding the weak repressor domain from BIN1 decreased activation and increased repression compared to FOXO3 alone (**Fig. S2A**, **S3A**).

While in general we found that strong repressors overpowered virtually any non-repressor effector domain they were paired with, we did find a few examples of activators that could prevent dual-functional domains from repressing strong promoters (**Fig. 4A**). These activators include the activator KRAB from ZN597, a variant KRAB domain that functions as an activator; the SH3 domain from ANM2 previously described in this work; and the activating SH3 domain from BTK. While BTK’s SH3 domain is known to regulate BTK kinase activity upon autophosphorylation^51^, and while BTK can translocate to the nucleus^52^, we are unaware of a previously described role for the SH3 domain in regulating gene expression of BTK targets.

We were excited to find a number of non-hit domains that could not alter gene expression when paired with negative controls but did affect the function of other effector domains (gray labels, **Fig. 4A**, **Fig. S5A**). For example, we found that the aforementioned domain from DPY30, a core subunit of the SET1/MLL methyltransferase complex that interacts with ASH2L to establish H3K4me3^53^, was able to ablate repressor function when paired with not only dual-functional domains but even weak to moderate strength repressors. Additionally, the N-terminal domain of DPF1, which links the NF-κB RelA/p52 heterodimer with SWI/SNF complex subunits to drive transcription^54^, was able to prevent repressor function from strong repressors, such as HERC2’s Cyt-b5 domain. We also found non-hit domains that prevented activation when paired with other effectors, including the a tile from the N-terminal disordered region of DNMT3B thought to be part of a broader region mediating interactions with the methyltransferases DNMT1 and DNMT3A^55^. The C2HR domain from the zinc finger ZNF496, which has been shown to overpower the variant activator KRAB present on the same transcription factor^22^, was similarly able to prevent all activators and all but the strongest dual-functional domains from driving gene expression. Altogether, these effectors did not themselves activate or repress transcription when paired with negative controls, but did modulate the activity of partner effectors on the same molecule in a manner consistent with the function of their native proteins.

We wondered how the distribution of effector combinations along both repression and activation log_2_(ON:OFF) scores varied between repressor-dual, activator-dual, and repressor-activator combinations. We found that while most repressor-dual combinations functioned as pure repressors, some dual-functional domains were able to combine with repressors to produce overall dual-functional combinations (**Fig. 4F**). In contrast, combining activators with repressors produced a larger number of effector pairs that neither activated the weak promoter nor repressed the strong promoter, with relatively fewer dual-functional combinations (**Fig. 4G**). Pairing pure activators with dual-functional domains produced combinations that mostly acted as dual activator-repressors, with a smaller fraction of combinations acting as pure activators without maintaining the repressor effect of the dual-functional domain (**Fig. 4H**).

### Systematic characterization of domains that influence KRAB-mediated repression

Our screen data so far indicated that virtually any concatenation including the ZNF10 KRAB domain functioned as a strong repressor (~180 KRAB-containing pairs). We were interested in investigating this KRAB domain in more detail, as it is widely used in the well-known CRISPRi system^56^, has been harnessed in conjunction with DNA methyltransferases to produce more durable epigenetic silencing^9,25^, and has been deeply characterized^21^. We decided to test whether this pattern of KRAB dominance held when using a larger panel of partner domains, and generated a lentiviral library encoding a set of concatenations fusing ZNF10 KRAB with a library comprised of ~5000 80-amino acid proteins sequences consisting of Pfam annotated domains from nuclearly localized proteins and a set of negative controls^21^. We transduced this library into K562 cells expressing a reporter gene driven by the strong pEF promoter (**Fig. 5A**) and recruited the concatenations to the reporter gene for 5 days by adding doxycycline. We used a lower doxycycline concentration than in our previous screens (100 ng/ml vs 1000 ng/ml); to slow down KRAB silencing (**Fig. S6A**) and thus allow for a wider dynamic range for measuring both decrease and increase of function. We measured the enrichment of domain pairs in the ON versus OFF populations (**Fig. S6B**) at day 2 of recruitment to determine the speed of silencing; we opted not to take a measurement on day 5 at the end of recruitment, as the population was virtually entirely silent (**Fig. 5B**).

**Figure 5.**
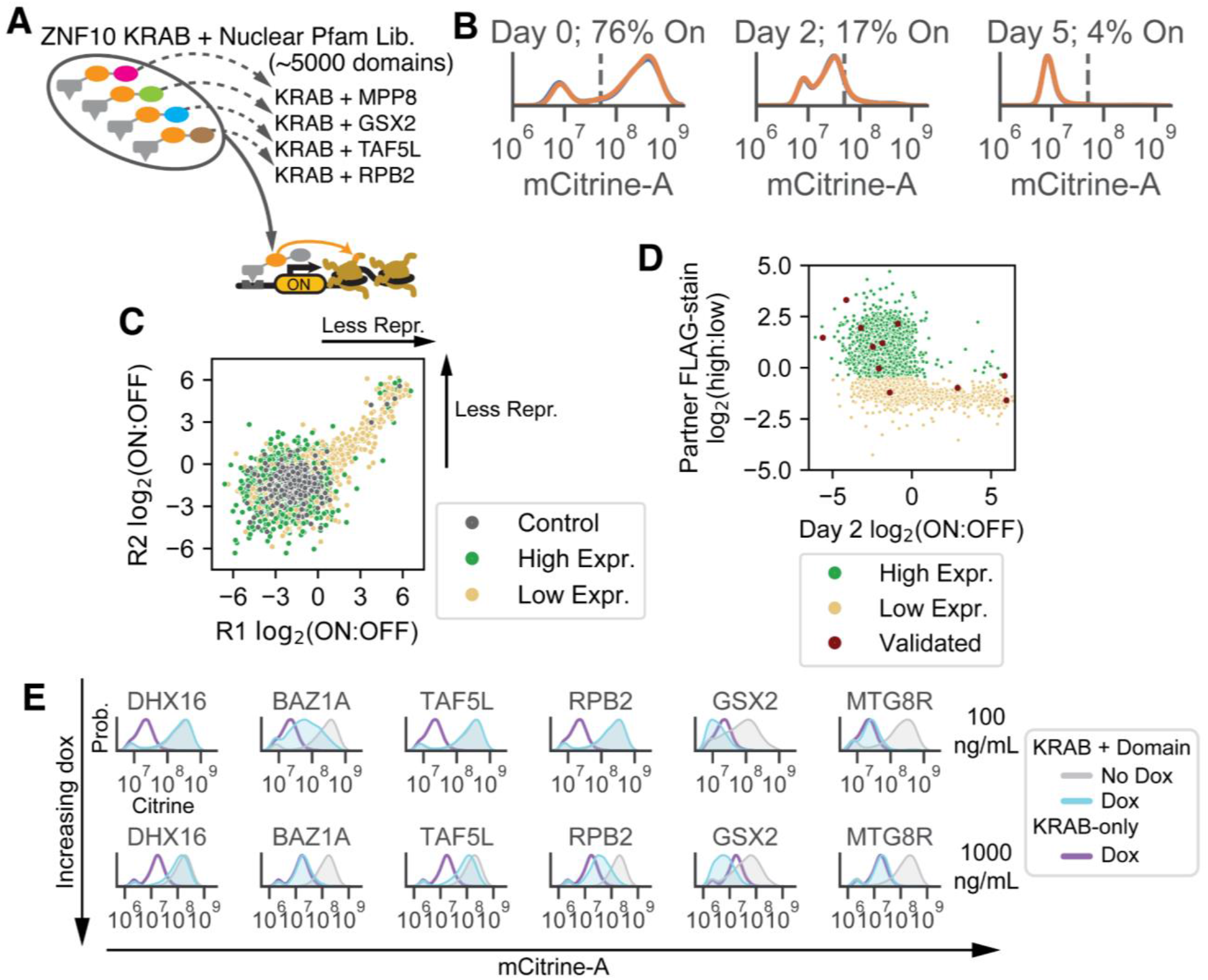
KRAB repression. A. Schematic of KRAB-Pfam screen. A library of ~5000 Pfam-annotated nuclear-localized domains^21^ was cloned into a backbone downstream of an rTetR DNA-binding domain fused to the ZNF10 KRAB repressor and recruited at the constitutive pEF promoter. B. Flow cytometry measurements of mCitrine fluorescence (x-axis) at 0, 2, and 5 days after 100ng/ml dox-mediated recruitment at the pEF promoter. Data shown consists of 2 replicates. C. Correlation between replicate 1 (x-axis) and replicate 2 (y-axis) of screen log_2_(ON:OFF) scores after 2 days of dox recruitment for KRAB-domain pairs where domain pairs were previously annotated as stable (green), unstable (yellow), or negative controls (gray). D. Correlation between screen log_2_(ON:OFF) scores (x-axis) and prior FLAG-stain measurements of KRAB partner domain expression (y-axis) for high-expressed domains (green), low-expressed domains (yellow). Domains that were individually validated via flow cytometry are shown in red. E. Probability density curves of mCitrine fluorescence for select domains that were individually validated using flow cytometry. Data shown is after 2 days of recruitment at 100 ng/mL dox (top row) or 1000 ng/mL dox (bottom row) at the pEF promoter. No dox controls are shown in gray, while recruitment of KRAB alone is shown in purple.

On day 2, the majority of pairs featuring well-expressed domains (as measured by FLAG staining before^21^) scored similarly to concatenations of KRAB with a negative control domain (**Fig. 5C-D**), consistent with the flow cytometry results showing that the majority of the cells were silenced (**Fig. 5B**). This matched our expectations from the prior screens, where concatenations featuring KRAB silenced similarly to KRAB on its own or with negative control domains. In this case, the negative controls consisted of a large set of random sequences and DMD fragments ^21^, and contained both well-expressed and poorly expressed proteins. We found that a number of domains that were lowly expressed on their own ablated KRAB function when paired with it (**Fig. 5C-D**). For example, we verified that the poorly expressed DHX16 OB_NTP and BAZ1A DDT domains inhibited KRAB function at 100 ng/mL dox (**Fig. 5E**, top). Interestingly, increasing the dox concentration to 1000 ng/mL dox permitted some silencing for KRAB-DHX16 and full silencing for KRAB-BAZ1A (**Fig. 5E**, bottom), consistent with the loss of function of these KRAB fusions coming from decreased protein abundance.

We also observed loss of KRAB function when fused to certain well-expressed domains from proteins that are part of the basic transcriptional machinery, namely the 2nd WD40 domain from TAF5L and the fork domain of RPB2. Consistent with the high-throughput measurements, individual validations at 100 ng/ml dox show a complete loss of KRAB silencing for the fusions with these domains (**Fig. 5E**, top row). The annotated RBP2 domain is smaller than the 80 aa sequence we used in our screen, and the 64 aa trimmed version had a lower capability of opposing KRAB, allowing for more silencing (**Fig. S6E**). Some amount of silencing was also restored at saturating dox concentrations for both KRAB-TAF5L and KRAB-RBP2 (**Fig. 5E**, bottom row), showing that KRAB can still dominate if enough of it is recruited at the locus. These results were corroborated by an overall increase in the rate of gene silencing when KRAB concatenations were recruited at 1000 ng/mL dox as compared to recruitment at 100 ng/mL dox (**Fig. S6F**, p=0.0239, paired t-test).

In the high-throughput measurements we also identified a small number of domains that increased KRAB silencing (**Fig. 5C**, bottom left). We validated that the library tile containing the homedomain from GSX2 increases KRAB silencing at day 2 when recruited at 100 and 1000 ng/ml doxycycline (**Fig. 5E**, left), consistent with its behavior as a repressor when recruited on its own in our previous Pfam screen^21^. Interestingly, the trimmed version that contains only the annotated homeodomain does not enhance KRAB silencing (**Fig. S6C**). In contrast, the NHR2 domain from MTG8R, on its own a slightly weaker repressor than GSX2’s homeodomain in our prior screens, was unable to modify KRAB-mediated silencing at both 100 ng/mL and 1000 ng/mL dox (**Fig. 5E**).

### Composing effector domains to generate multifunctional synthetic transcription factors

We wanted to take advantage of the fact that the KRAB repressor is dominant over activators to build a more versatile transcription factor that can switch between repressor and activator based on addition or removal of a second drug. We engineered a version of our rTetR-FOXO3-ZNF10 KRAB concatenation plasmid where the two effector domains were separated by a StaPL domain, which cleaves itself in the absence of asunaprevir (ASV)^57^ (**Fig. 6A**). Thus, in the absence of ASV, the KRAB domain would be cleaved, leaving behind the dual-functional rTetR-FOXO3, while upon addition of the ASV inhibitor, the KRAB domain would act as a dominant repressor (**Fig. 6B**).

**Figure 6.**
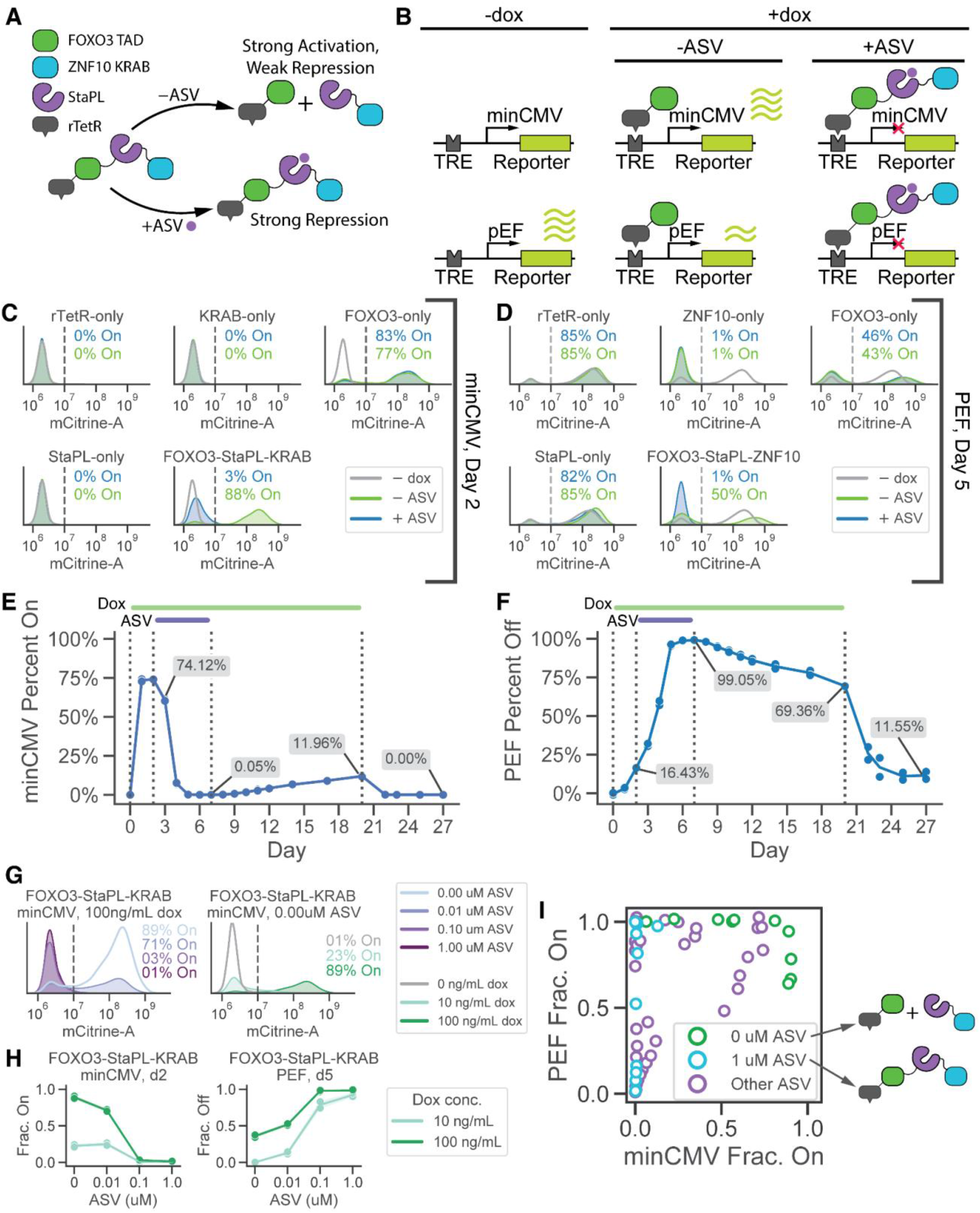
A synthetic TF that can switch from strong repressor to strong activator. A. Schematic of synthetic transcription factor system. An ASV-inducible StaPL (purple) is fused in between FOXO3’s TAD (green) and ZNF10’s KRAB (blue) domains. In the absence of ASV, the StaPL and KRAB are cleaved off, resulting in only FOXO3’s TAD being recruited to the reporter. Adding ASV stabilizes the StaPL domain, resulting in recruitment of FOXO3’s TAD fused to ZNF10’s KRAB domain. B. Expected dynamics of gene expression at the minCMV promoter (top row) or pEF promoter (bottom row) without dox (left), with dox and no ASV (middle), and dox and ASV (right). C. Histograms of transcriptional activation after 2 days of recruitment to the synthetic reporter driven by the minCMV promoter in the presence (blue) or absence (green) of ASV, with the 0 ng/mL dox control shown in gray. D. Histograms of transcriptional repression after 5 days of recruitment to the synthetic reporter driven by the pEF promoter in the presence (blue) or absence (green) of ASV, with the 0 ng/mL dox control shown in gray. E. Transcriptional activation after ASV washout. K562 cells expressing the minCMV-driven reporter and the FOXO3-StaPL-KRAB construct were treated with doxycycline during days 0-20 (green line) and ASV during days 2-7 (purple line). Cells were washed 1x with PBS when removing ASV and doxycycline. F. Transcriptional repression after ASV washout. K562 cells expressing the pEF-driven reporter and the FOXO3-StaPL-KRAB construct were treated with doxycycline during days 0-20 (green line) and ASV during days 2-7 (purple line). Cells were washed 1x with PBS when removing ASV and doxycycline. G. Histograms of transcriptional activation after 2 days of recruitment to the synthetic reporter driven by the minCMV promoter with either 100 ng/mL dox and varying doses of ASV (purple, left) or 0 uM ASV and varying doses of dox (green, right). H. Fraction of cells activated after 2 days of recruitment to the synthetic reporter driven by the minCMV promoter (y-axis) for different ASV doses (x-axis), and for varying doses of doxycycline (light vs dark green). I. Scatter of minCMV-reporter fraction on (x-axis) versus pEF-reporter fraction on (y-axis) measured across all FOXO3-StaPL-KRAB experiments. Each point constitutes 1 timepoint, and the color of each point indicates the ASV dose.

We first verified that the construct worked as expected when driving gene activation at the minCMV promoter (**Fig. 6C**) and gene repression at the pEF promoter (**Fig. 6D**, **Fig. S7A**) at maximum recruitment with a saturating dose of dox (1000 ng/mL). At minCMV, the rTetR-FOXO3-StaPL-KRAB drove strong gene activation in the absence of ASV (**Fig. 6C**, bottom right green) to a similar level as rTetR-FOXO3 only (**Fig. 6C**, top right). At the same promoter, addition of ASV reduced activation by the rTetR-FOXO3-StaPL-KRAB to a minimum (**Fig. 6C**, bottom right blue), comparable to rTetR-KRAB recruitment, and consistent with KRAB dominating over FOXO3. At pEF, the rTetR-FOXO3-StaPL-KRAB repressed virtually all cells upon ASV addition (**Fig. 6D**, bottom right blue), the same as rTetR-KRAB alone. Without ASV, this fusion still repressed 50% of the cells (**Fig. 6D**, bottom right green), consistent with rTetR-FOXO3 alone being able to silence pEF in about ~54-57% percent of cells (**Fig. 6D**, top right). As expected, the negative controls rTetR alone and rTetR-StaPL did not change gene expression at either promoter (**Fig. 6C-D**). Altogether, the construct behaved as expected: adding ASV changed its behavior from FOXO3-like to KRAB-like.

The ability to toggle the FOXO3-StaPL-KRAB TF from activator to repressor and then back to activator again allowed us to test whether the activator behaves differently at a promoter before and after KRAB-induced repression. In order to do this, we added dox to cells containing this fusion for a period of 20 days to induce recruitment, and within this interval we varied ASV to toggle between FOXO3 (-ASV) and KRAB dominant (+ASV) (**Fig. 6E-F**). At the minCMV reporter, we saw rapid gene activation upon addition of doxycycline that was lost when ASV was added and KRAB recruited to the promoter (**Fig. 6E**). Removal of ASV, leading to recruitment of FOXO3 alone at this KRAB-silenced minCMV promoter produced much slower gene activation than at the unsilenced minCMV promoter (**Fig. 6E**, days 0-2 vs. 7-20). The slow reactivation upon ASV washout suggests that while FOXO3 may be able to drive gene expression from a minimal promoter at a permissive locus, it is less efficient at reactivating that promoter after it is silenced with potentially methylated and/or compacted chromatin. At the pEF promoter (**Fig. 6F**), we observed a slow reduction in gene expression without ASV during days 0-2, consistent with the dual FOXO3 acting as a weak repressor at pEF. The percentage of cells silenced rapidly increased upon the addition of ASV and recruitment of KRAB culminating in complete silencing by day 7. After removal of ASV, we saw slow but measurable reactivation that increased dramatically upon doxycycline removal (**Fig. 6F**, days 7-20 vs. 20-27), suggesting that while FOXO3 can behave as an activator and drive gene expression at minCMV, its repressive capacity may inhibit proper reactivation of the pEF promoter.

In order to characterize the range of behaviors achievable with this inducible synthetic TF, we varied the dosing of both ASV and doxycycline while recruiting the rTetR-FOXO3-StaPL-KRAB construct at both promoters. At 100 ng/mL dox increasing doses of ASV led to decreased gene activation at the minCMV promoter (**Fig. 6G** left, **S7B**), consistent with the expectation that increasing doses of ASV led to an increase in the species containing KRAB relative to FOXO3 only. We also found that increasing the dose of dox at 0 ASV led to increasing gene activation at the minCMV promoter, consistent with a model in which the absence of ASV leads to virtually minimal KRAB recruitment (**Fig. 6G** right, **Fig. S7B**). Low to moderate doses of ASV, 0.01uM and 0.1uM, produced intermediate profiles between FOXO3 and KRAB (**Fig. S7B-C**). Altogether, these results suggest a model where increasing the dose of ASV titrates between FOXO3-like and KRAB-like behavior, while increasing the dose of dox increases the degree of gene activation and/or repression.

We wished to understand what new profiles of gene expression the combined FOXO3-StaPL-KRAB transcription factor could generate that neither FOXO3 nor KRAB could produce on their own. For each experiment performed with the transcription factor, we plotted for each timepoint the fraction of cells actively expressing the pEF reporter versus the fraction expressing the minCMV promoter (**Fig. 6I**). We found a large number of states that the synthetic inducible TF was able to access at intermediate ASV doses (**Fig. 6I**, purple) that could not be achieved at 0uM ASV (FOXO3-like behavior) or 1uM ASV (KRAB-like behavior). These states were principally composed of situations where both minCMV and pEF were expressed in only a fraction of cells; FOXO3-like behavior mainly produced high levels of cells with minCMV on (**Fig. 6I**, green), while KRAB-like behavior did not permit minCMV expression at all (**Fig. 6I**, blue). In conclusion, we found that composing FOXO3 and KRAB in this manner and varying the dose of ASV produced distinct profiles of gene expression not achievable with either effector domain individually.

## Discussion

Despite considerable efforts to parse the combinatorial logic of TF binding and gene regulation^19,58–60^, our understanding of how multiple distinct functional transcriptional effector domains work together within a single TF remains limited. Improved characterization of the combined function of effector domains is important for understanding the function of natural TFs featuring multiple effector domains^2^, for building complex synthetic biology tools to manipulate gene expression^61^, and for building cell therapies to detect and treat diseases^26,62–65^. Here, we present the results of screening thousands of effector domain combinations, another step towards uncovering the basic principles of combinatorial gene regulation.

We found that weak activators can synergize to drive robust gene expression even when the individual activators being paired were not particularly strong. This is in agreement with previous results showing that multiple activation domains can act synergistically in yeast when recruited at a synthetic reporter^7,14^, and in human cells when recruited at reporters or endogenous genes using dCas9^24,66^. At the other extreme, we found that some of the strongest activators, which in our system acted as dual-functional domains that could also repress a constitutive promoter, were often antagonistic: when paired with each other, they produced less gene activation from a minimal promoter than when recruited on their own. Antagonism between transcriptional activators has been reported in other contexts: TFs have been shown to interfere with each other’s ability to activate genes via direct interactions with each other^67^ or via squelching mechanisms involving the sequestration of coactivator proteins^68^. However, our findings suggest a more general negative feedback mechanism triggered by high levels of activators at a promoter; more thorough confirmation of this phenomenon and subsequent investigations into its molecular underpinnings are still needed.

Our activation data showed a tight coupling between the fraction of actively transcribing cells and the degree of transcription in those cells, consistent with manipulation of transcriptional bursting dynamics^49,69^). Recently, certain TFs were shown to increase only burst size or only burst frequency, depending on their molecular mechanism of action^70^. While our long-lived protein reporter does not allow us to differentiate between changes in burst size or frequency for our combinations, it would be interesting to repeat these high-throughput measurements coupled with RNA FISH or a destabilized reporter that allows extraction of burst parameters.

We found that repressors generally overpowered activators, although select activator domains could weaken the repressor function of dual-functional or weakly repressive domains. We confirmed this result more thoroughly via an in-depth screen measuring KRAB function when paired with a panel of 5,000 domains from nuclearly localized proteins, and found that KRAB maintained its repressive function when paired with most domains except ones that had decreased expression and a few domains from general TAFs and RNA polymerase. We used these insights to build a synthetic transcription factor whose function could be switched from a dual-functional FOXO3-like profile (that activates a minimal promoter and weakly represses a constitutive one) to a repressive KRAB-like profile via the addition of the small molecule drug ASV. We used this switchable TF to show that both the activation and repression functions of the FOXO3 effector domain changed when the target promoter was first silenced by KRAB. While activation from a minimal promoter was impaired by prior silencing as expected, recruitment of FOXO3 increased KRAB-mediated epigenetic memory and reduced reporter gene reactivation at the pEF promoter. Importantly, since ASV tuned the behavior of this synthetic TF independent of dox-mediated recruitment, we could generate gene expression profiles that were not accessible with either individual effector domain alone. Looking forward, this synthetic switchable TF can be used to split a cell population into well-defined percentages that express the desired combination of two target genes, e.g., 50% of the cells expressing gene 1 and 50% expressing gene 2. Such flexibility could allow for more sophisticated and complex gene circuits and the engineering of “higher-order” cell behaviors and programs^71^.

When we started this study, we had a limited number of validated activation and repressive domains to test in combination. Recently, however, we have identified a much larger set of effector domains from human TFs^20^. Prior efforts have characterized individual examples of multiple effector domains being combined within a larger TF^22,72^; it would be instructive to design a library that systematically tests combinations of domains that come from the same TF and compare these results with recent ORFeome screens measuring activation and repression of full-length TFs^33^. Moreover, emerging work has begun to connect TF and effector function with measurements of their affinities for transcriptional coactivators and corepressors^13,33,73^. In addition, while our work characterized effector combinations at two promoters in one human cell line, we are excited for future efforts to broaden these settings to include other DNA-binding domains beyond rTetR, other promoters, and other cell types. Such efforts will undoubtedly aid projects aiming to build robust tools for cell engineering that can function across multiple contexts^74–76^.

## Supporting information

Supplementary Table 1 -- Co-Recruit_Oligo_Library

Supplementary Table 2 -- Co-Recruit_Screen_Data

Supplementary Table 3 -- KRAB_Pfam_Screen_Data

---

**Figure S1.**
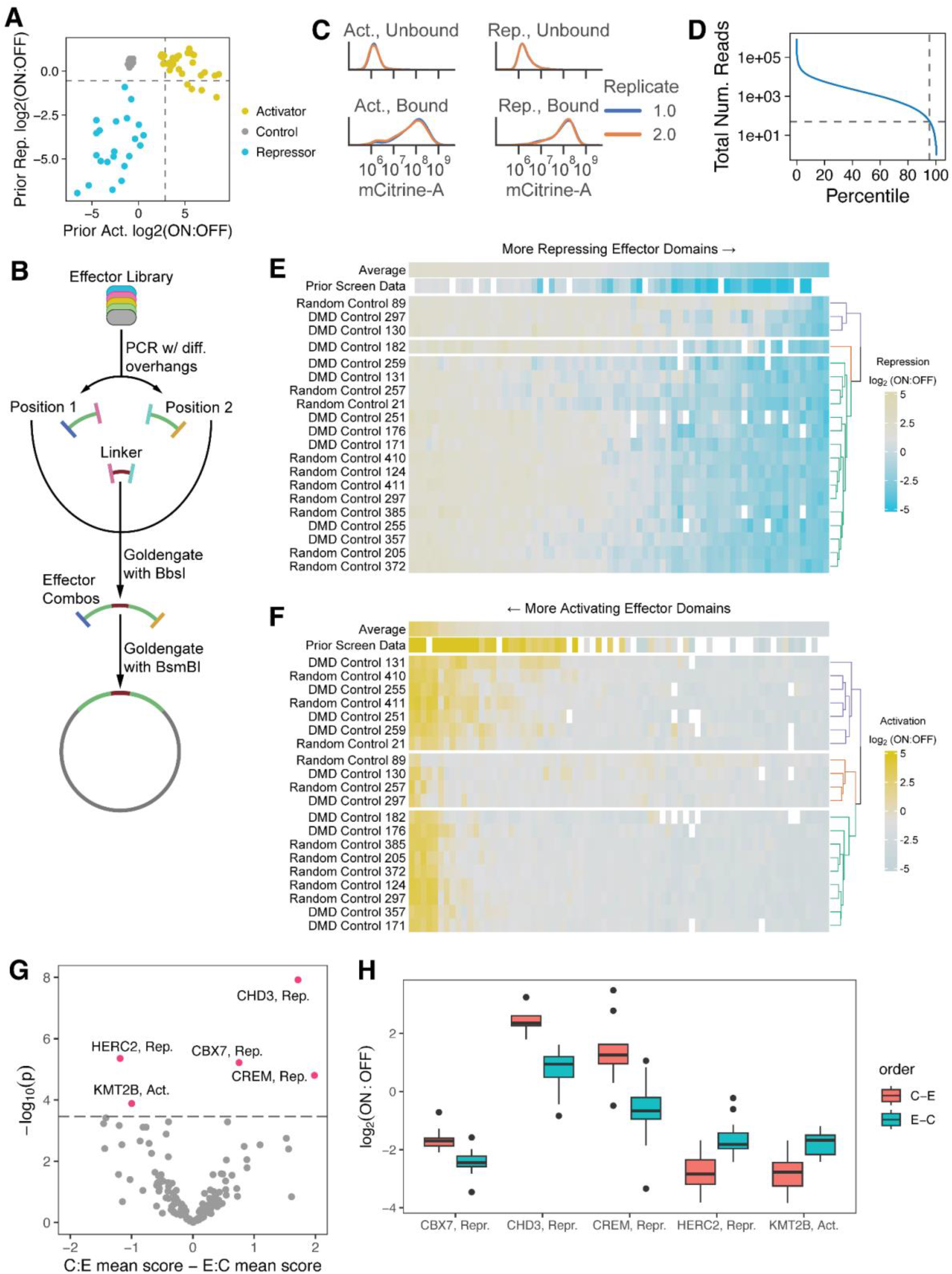
Screen Quality Control and Filtering. A. Scatterplot of prior data^21^ for both activation at minCMV (x-axis) and repression at pEF (y-axis) for individual domains chosen in the screen, colored based on their prior behavior. Controls are random 80AA sequences or tiles from the DMD protein. Dashed lines represent hit thresholds in the prior screen. B. Schematic of co-recruit cloning strategy. Domain library is PCRed with distinct overhangs to produce position 1 and position 2 libraries, which are then joined to the XTEN linker in one Goldengate reaction to produce a library of effector concatenations. Concatenations are then ligated into a lentiviral backbone via a second Goldengate reaction. C. Probability density (y-axis) of mCitrine fluorescence levels (x-axis) for bound (top) and bound (bottom) cells post-magnetic separation for activation (left) and repression (right) screens, with each replicate shown in a different color. D. Total number of reads for each domain pair (y-axis); domain pairs are ordered from most to least reads, with x-axis marks indicating what percentile of pairs had more reads than those at that x-value. E. Clustermap of controls (rows) when paired with each repressing effector domain (columns), with average effector domain profiles shown along the top. Color indicates the repression log_2_(ON:OFF) score for a particular control-effector pair. Domains filtered out were random control 89 and DMD controls 130, 182, and 297. F. Clustermap of controls (rows) when paired with each activating effector domain (columns), with average effector domain profiles shown along the top. Color indicates the activation log_2_(ON:OFF) score for a particular control-effector pair. Domains filtered out were random controls 89 and 257 and DMD controls 130 and 297. G. Comparison of orientations for each domain. Difference between the average score with the effector C-terminal to the control domain (C:E) and the average score with the effector N terminal to the control domain (E:C) (x-axis) is plotted against the negative log of the p-value as determined by a Welch’s t-test. Dashed line indicates the Bonferroni-corrected significance threshold, and domains with significant differences between orientations are marked in red. H. For domains with statistically significant orientation-dependence, box-and-whisker plots of their log_2_(ON:OFF) scores are shown for each of the 2 possible orientations. Orientations that weakened the domains’ effects were removed from downstream analysis.

**Figure S2.**
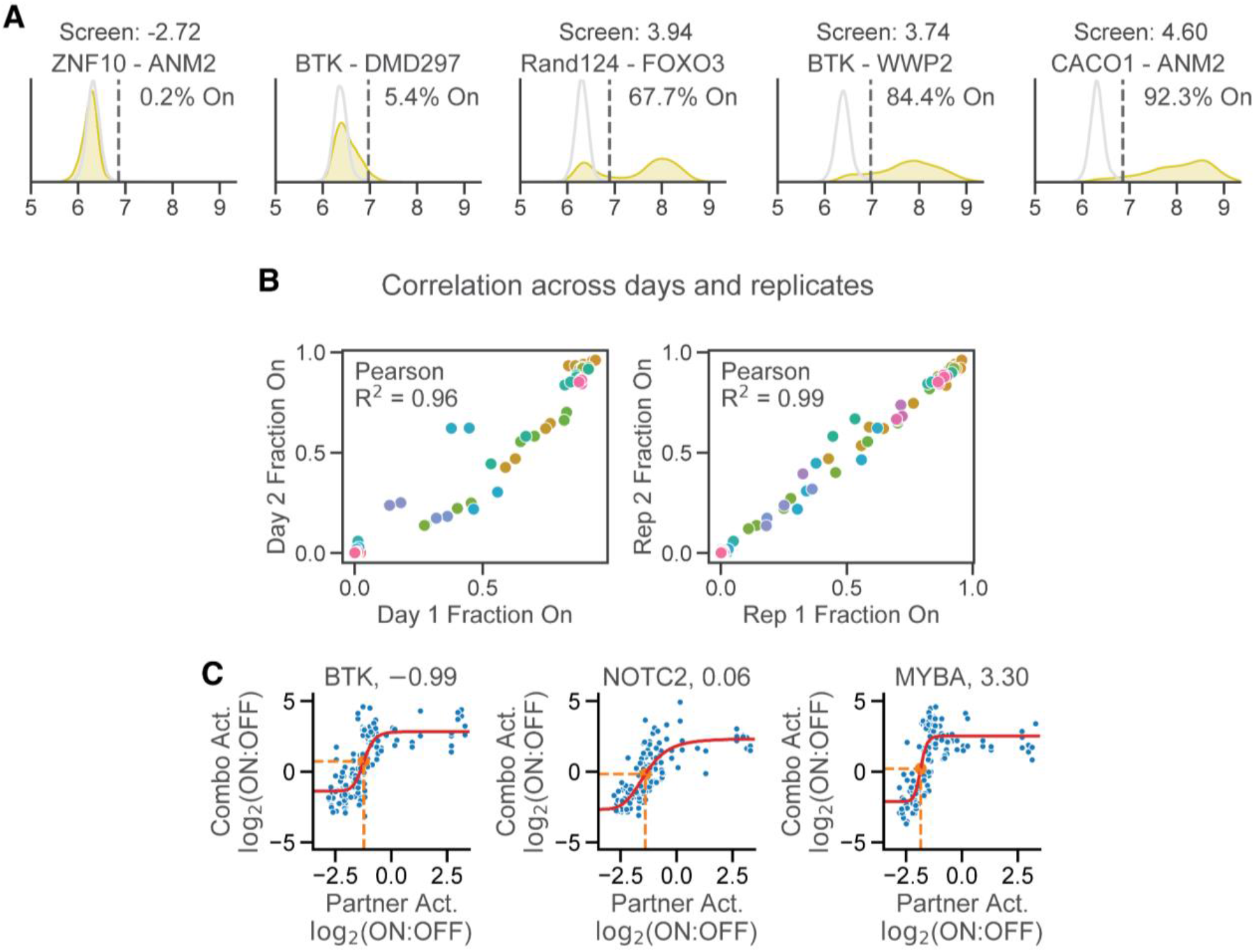
Activator Validations and Modeling. A. Example low-throughput validations of activator-negative control and activator-activator pairs. No doxycycline recruitment is shown in light gray, while the histogram of fluorescence after 2 days of dox recruitment is shown in yellow. Dashed lines represent the fluorescence threshold for calling cells On in reporter expression. B. Correlations between days 1 and 2 (left) and replicates 1 and 2 (right) for measuring the activation strength of domain concatenations. Colors indicate different constructs. C. Sigmoidal shape of activator behavior in the high-throughput screen. For BTK (left), NOTC2 (middle), and FOXO3 (right), the control-paired score of its partner in any given pair (x-axis) is plotted against the score of the combination of the domain and that partner (y-axis) for all concatenations featuring the given domain.

**Figure S3.**
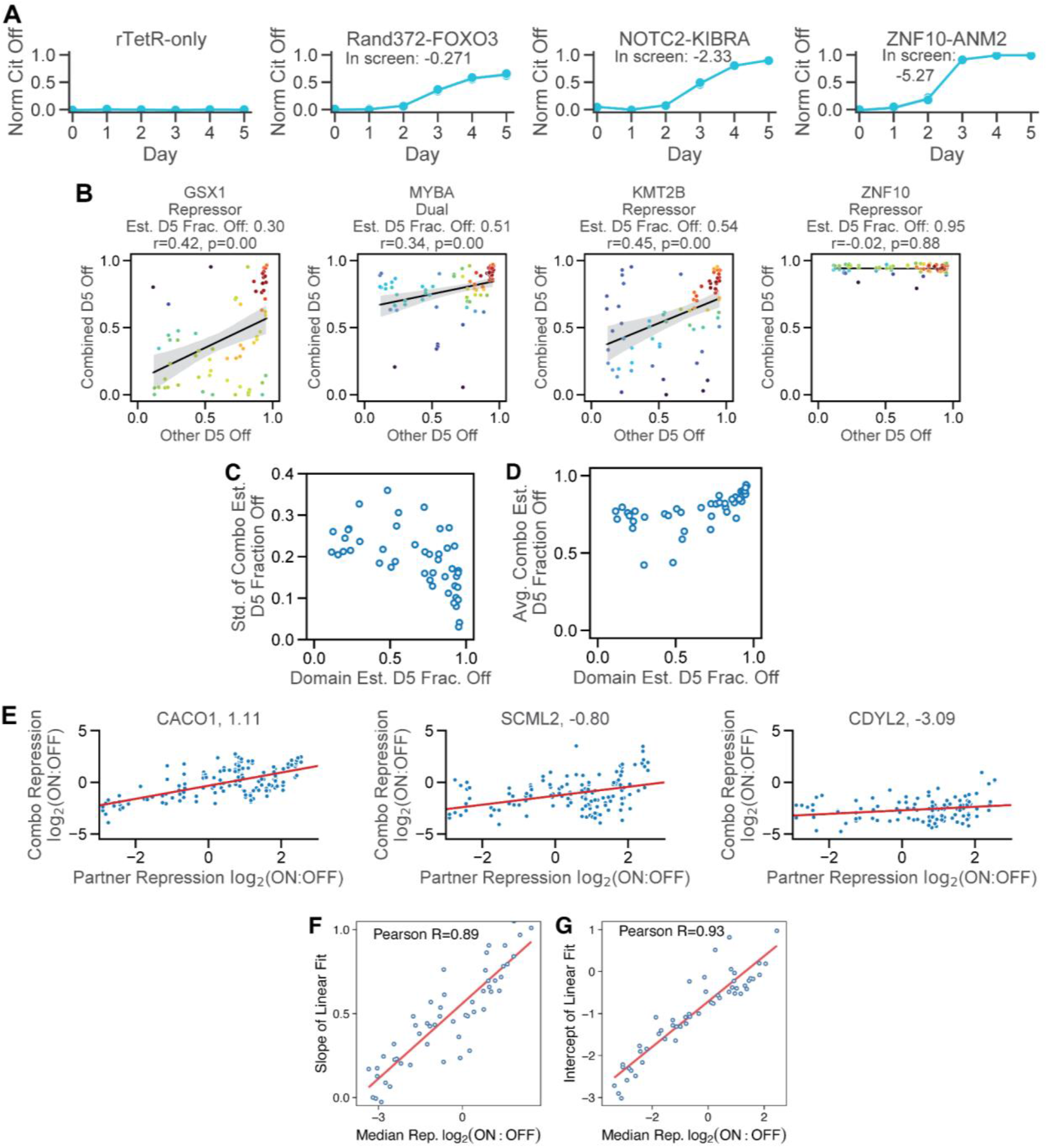
Repressor validations and screen data analysis. A. Low-throughput measurement of repressor function of various domain combinations, with Rand372 denoting a negative control domain. Cells were labeled as Off when their mCitrine-A fluorescence dropped below 10^7^ (**Fig. 3B**). B. Effect of partner estimated day 5 fraction off on combination day 5 fraction off for repressor domains. For each repressor or dual domain (GSX1, MYBA, KMT2B, or ZNF10) the estimated day 5 fraction off of a given partner of that domain (x-axis) is plotted against the estimated day 5 fraction off for the concatenation (y-axis). Color denotes density as computed by a kernel density estimator, with red constituting denser, and blue less dense. C. Estimated day 5 fraction off (x-axis) for each domain versus the standard deviation of the estimated day 5 fractions off for all combinations including that domain (y-axis) for all repressor-repressor, repressor-dual, and dual-dual pairs. D. Estimated day 5 fraction off (x-axis) for each domain versus the average of the estimated day 5 fractions off for all combinations including that domain (y-axis) for all repressor-repressor, repressor-dual, and dual-dual pairs. E. Linear shape of repressor behavior in the high-throughput screen. For a given domain (CACO1 - left, SCML2 - middle, CDYL2 - right), the control-paired score of its partner in any given pair (x-axis) is plotted against the score of the combination of the domain and that partner (y-axis) for all concatenations featuring the given domain. Red lines indicate linear fits to the data. F. For all repressor domains, the control-paired strength of the repressor (x-axis) is plotted against the slope of the corresponding linear fit to screen data (y-axis). Line shown is the line of best fit. G. For all repressor domains, the control-paired strength of the repressor (x-axis) is plotted against the y-intercept of the corresponding linear fit to screen data (y-axis). Line shown is the line of best fit.

**Figure S4.**
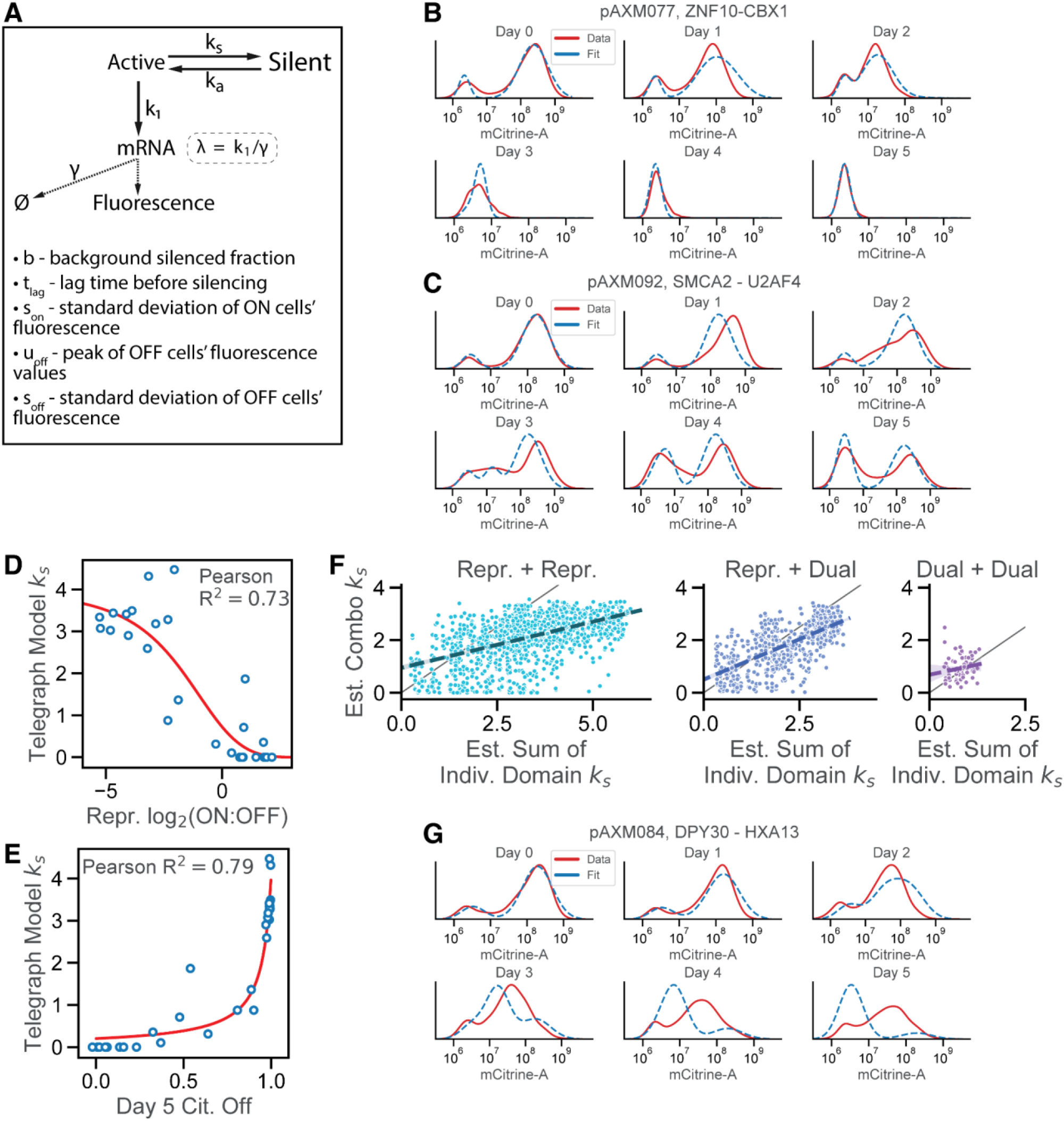
Repressor modeling. A. Brief description of the silencing model (**Materials & Methods**). Cells in the Active state produce mRNA at rate k1, which decays with rate γ. We define λ = k_1_ /γ to be the average amount of mRNA in cells at steady state when Active. Cells in the Active state are silenced with rate k_s_ and reactivate at rate k_a_ upon recruitment. The model incorporates a permanently silent population *b* of background silenced cells. We assume a lag time tlag before silencing. The standard deviation of the Active cells fluorescence is denoted s_on_ while the standard deviation of the Silent cells fluorescence is s_of_. The fluorescence peak of Silent cells is defined as u_off_. B. Example fit of the silencing model for ZNF10-CBX1, with fluorescence indicated on the x-axis and density on the y-axis. Data gathered from flow cytometry measurements is shown in red, while the fitted model is shown in blue for each day (0-5). C. Example fit of the silencing model for SMCA2-U2AF4, with fluorescence indicated on the x-axis and density on the y-axis. Data gathered from flow cytometry measurements is shown in red, while the fitted model is shown in blue for each day (0-5). D. Correlation between repression log_2_(ON:OFF) scores for validated concatenations (x-axis) and k_s_ values as estimated by the silencing model (y-axis), with best-fit sigmoid shown in red. E. Correlation between day 5 fractions of cells silenced for validated concatenations (x-axis) and k_s_ values as estimated by the silencing model (y-axis), with best-fit asymptotically increasing curve shown in red. F. For repressor-repressor (left), repressor-dual (middle), and dual-dual (right) pairs, the sum of the estimated k_s_ for both individual domains (x-axis) is plotted against the estimated k_s_ for the combination (y-axis). Line shown is a best-fit linear regression with a 95% confidence interval shaded in. Gray line indicates the y=x line. G. Example fit of the silencing model for DPY30-HXA13, with fluorescence indicated on the x-axis and density on the y-axis. Data gathered from flow cytometry measurements is shown in red, while the fitted model is shown in blue for each day (0-5).

**Figure S5.**
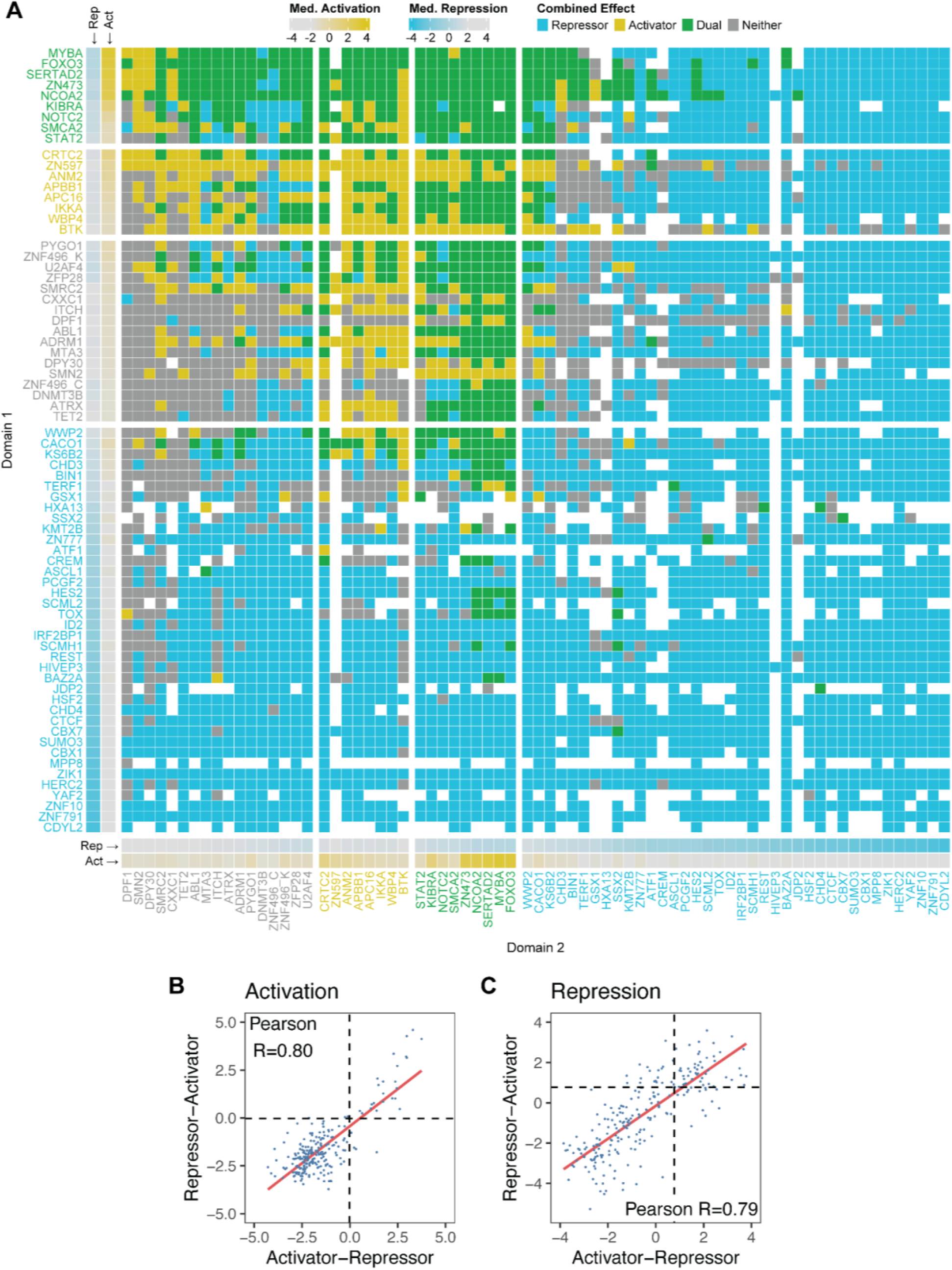
Activator/repressor interactions and orientation dependence. A. Representation of effector-effector interactions measured by the high-throughput screen. Domains in position 1 are arrayed along the y-axis separated into 4 categories (from bottom to top: repressors, non-hits, activators, and dual-functional domains), with the stronger duals, activators, and non-hits as measured by control-paired activation scores on the top of their respective groups. Repressors, in contrast, are ordered by their repression score, with the strongest repressors at the bottom. Boxes just to the right of each activator domain indicate the control-paired activation/repression score for each effector. Domains in position 2 are arrayed along the x-axis separated into 4 categories (from left to right: non-hits, activators, dual-functional domains, and repressors), with the stronger repressors within each category as measured by control-paired activation scores on the right. Boxes just above each repressor domain indicate the control-paired activation/repression score for each effector. Boxes in the interior represent combinations of the corresponding activator and repressor, and are colored based on whether the combination acts as a repressor only (blue), activator only (yellow), both a repressor and an activator (green), or neither (gray). B. Correlation of activator-repressor pair activation log_2_(ON:OFF) scores when recruited at minCMV (to test activation) between an orientation where the activator is N-terminal to the repressor (x-axis) and an orientation where the activator is C-terminal to the repressor (y-axis). Red line represents the line of best fit. Dashed lines indicate the threshold for an effector domain combination to be labeled an activator from **Fig. 1E**. C. Correlation of activator-repressor pair repression log_2_(ON:OFF) scores when recruited at pEF (to test repression) between an orientation where the activator is N-terminal to the repressor (x-axis) and an orientation where the activator is C-terminal to the repressor (y-axis). Red line represents the line of best fit. Dashed lines indicate the threshold for an effector domain combination to be labeled an activator from **Fig. 1F**.

**Figure S6.**
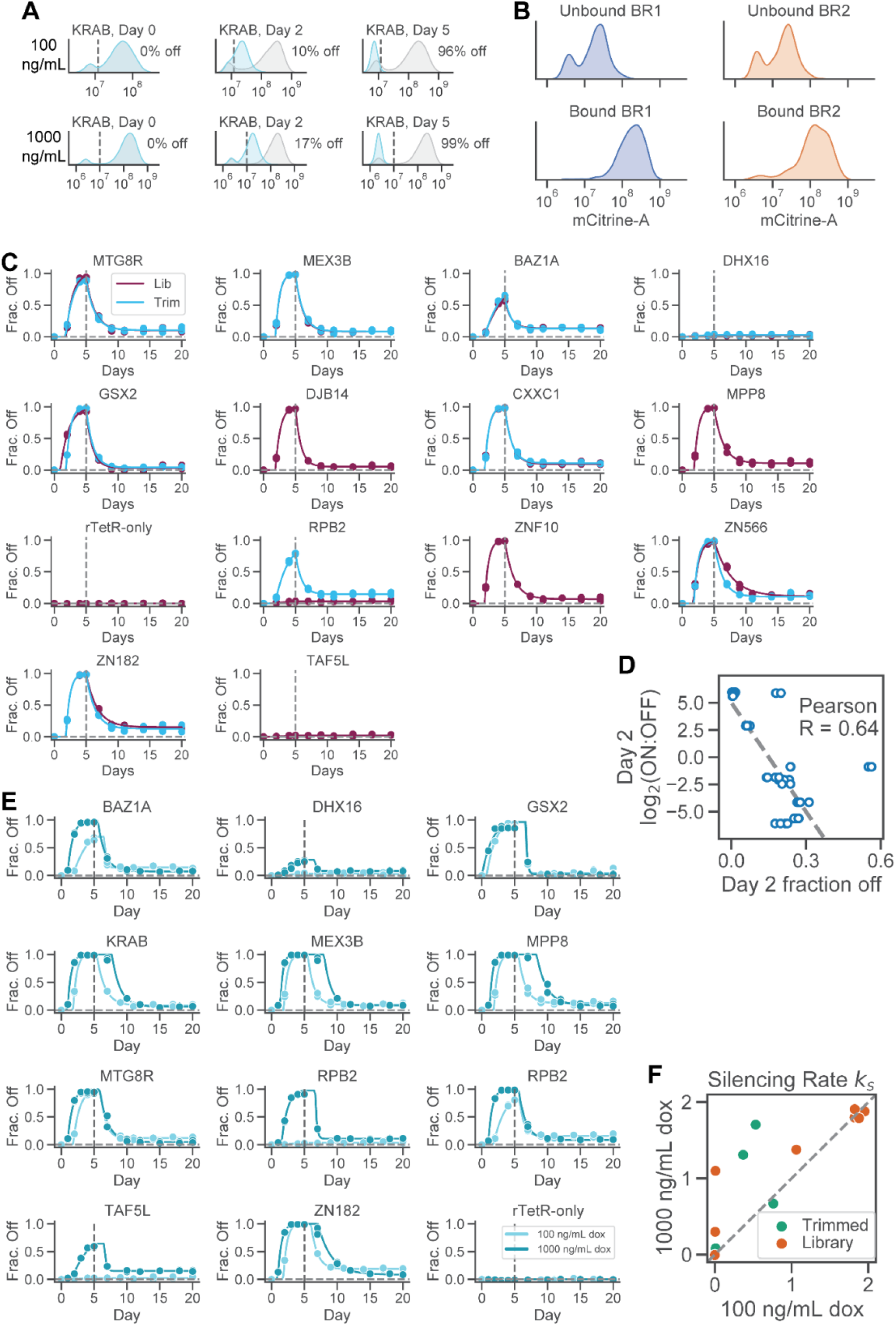
KRAB screen quality control and filtering. A. Probability density (y-axis) of mCitrine fluorescence levels (x-axis) for KRAB at days 0, 2, and 5 when recruited at either 100 ng/mL dox (top) or 1000 ng/mL dox (bottom). Vertical dashed lines indicate the threshold for labeling cells as ON or OFF. B. Probability density (y-axis) of mCitrine fluorescence levels (x-axis) for unbound (top) and bound (bottom) fractions post-magnetic separation for screen replicates 1 (left) and 2 (right) at day 2 after dox addition C. Individual recruitment of domains in low throughput to a reporter gene driven by the strong pEF promoter. Effectors in the library are composed of 80 amino acids surrounding the Pfam-annotated domain (lib, purple line) but were also tested in trimmed formats comprising only the annotated domain (trim, blue line). Doxycycline was added to cells for 5 days, and then cells were monitored for gene reactivation 20 days post-dox. Fractions of cells silenced were normalized with respect to a no-doxycycline control (**Materials and Methods**). Datapoints were fit using the 3-state phenomenological model^10,49^. D. Correlation between the fraction of cells silenced at day 2 of recruitment during low-throughput measurements (x-axis) and the corresponding log_2_(ON:OFF) scores from the KRAB-Pfam screen data (y-axis). Gray line shown is the line of best fit. E. Recruitment of domains at varying strength. KRAB-domain pairs were recruited to the pEF reporter in K562 cells at a dox concentration of either 100 ng/mL (light blue) or 1000 ng/mL (dark blue) for 5 days. Cells were monitored for 20 days after doxycycline release to measure gene reactivation. Fractions of cells silenced were normalized with respect to a no-doxycycline control. F. Stronger silencing at higher doxycycline doses. Rates of gene silencing computed using the 3-state model are compared for each domain measured in low-throughput at 100 ng/mL dox (x-axis) and 1000 ng/mL dox (y-axis). Gray line shown is y=x.

**Figure S7.**
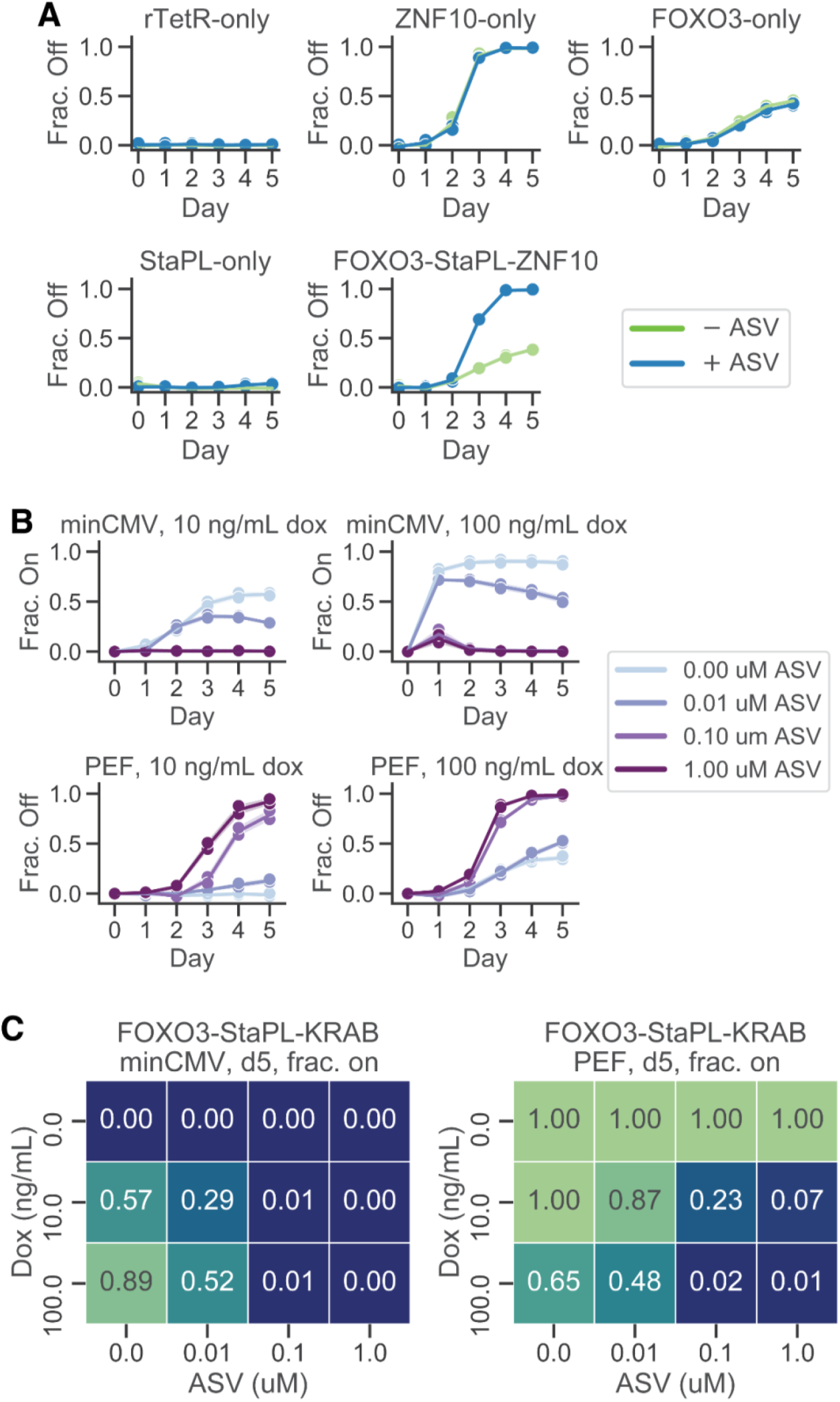
Functional characterization of FOXO3-StaPL-KRAB. A. Recruitment of rTetR-only, ZNF10-only, FOXO3-only, StaPL-only, and FOXO3-StaPL-KRAB to the pEF promoter at 1000 ng/mL dox for 5 days. Fraction of silent cells normalized to background (y-axis) is plotted against days of recruitment (x-axis). Data shown is the average of 2 replicates. B. Recruitment of FOXO3-StaPL-KRAB to the minCMV (top) or pEF (bottom) promoters for 5 days at different dox doses (100 ng/mL - left, 1000 ng/mL - right) and 4 different ASV doses. Each line shown is the average of 2 replicates. C. Fraction of cells activated at minCMV after 5 days of recruitment (left) and repressed at pEF after 5 days of recruitment (right) at 3 different doxycycline doses (rows) and 4 different ASV doses (columns). Data shown is the average of 2 replicates.

## Supplementary Tables

**Supplementary Table 1 – Base Oligo Library**

Sequences for domains used in the combinatorial recruitment screen related to Figs. 1-4 are attached in a CSV file.

**Supplementary Table 2 – Corecruit screen CSV**

Domain combinations and their corresponding measurements in the combinatorial recruitment screens for activation are attached in a CSV file.

**Supplementary Table 3 – KRAB+Pfam screen CSV**

Domains fused to KRAB and corresponding measurements in the repression screen related to Fig. 5 are attached in a CSV file.

## Materials & Methods

### Library design

Library members were chosen from the Nuclear Pfam library described in^21^. A total of 20 negative control domains, 10 randomers and 10 tiles of the DMD protein; 30 activator domains; and 50 repressor domains were chosen for cloning. Of the 50 repressors, 6 were discovered to have errors introduced during design, and were discarded from downstream analysis. All domains, including those eventually discarded, are listed in **Supplementary Table 1**. After library assembly, DNAChisel^77^ was used to optimize coding sequences by removing duplicates, 7xC homopolymers, BsmBI and BbsI restriction sites, and rare codons. Codon usage was matched to human codon prevalence and GC content was restricted to be between 20% and 75% in any 50-nucleotide window and between 25% and 65% globally.

### Cell culture

Cell culture was performed as described in^20^. Briefly, all experiments presented here were carried out in K562 cells (ATCC, CCL-243, female). Cells were cultured in a controlled humidified incubator at 37C and 5% CO2, in RPMI 1640 (Gibco, 11-875-119) media supplemented with 10% FBS (Omega Scientific, 20014T), and 1% Penicillin-Streptomycin-Glutamine (Gibco, 10378016). HEK293T-LentiX (Takara Bio, 632180, female) cells, used to produce lentivirus, as described below, were grown in DMEM (Gibco, 10569069) media supplemented with 10% FBS (Omega Scientific, 20014T) and 1% Penicillin Streptomycin Glutamine (Gibco, 10378016). minCMV and pEF reporter cell line generation is described in^21^. pEF and minCMV promoter reporter cell lines were generated by TALEN-mediated homology-directed repair to integrate donor constructs (pEF promoter: Addgene #161927, minCMV promoter: Addgene #161928) into the *AAVS1* locus by electroporation of K562 cells with 1000 ng of reporter donor plasmid and 500 ng of each TALEN-L (Addgene #35431) and TALEN-R (Addgene #35432) plasmid (targeting upstream and downstream the intended DNA cleavage site, respectively). After 7 days, the cells were treated with 1000 ng/mL puromycin antibiotic for 5 days to select for a population where the donor was stably integrated in the intended locus. Fluorescent reporter expression was measured by flow cytometry.

### Co-recruit cloning

Cloning for the combinatorial screen proceeded in two stages; in the first, domain-linker-domain concatenations were assembled, and in the second, concatenations were placed into a lentiviral backbone. Domains in the N-terminal and C-terminal position were synthesized as 2 separate oligonucleotide pools (Twist Biosciences). Each pool was PCR amplified in a clean PCR hood to avoid DNA contamination. Each pool was split into 6x 50 uL reactions that were PCR amplified for 21 cycles with 5 ng template, 1 uL of each 10mM primer, 1uL of Herculase II polymerase (Agilent), 1 uL of DMSO, 1 uL of 10mM dNTPs, and 10 uL of 5x Herculase buffer (Agilent). Reaction mixes were thermocycled at 98C for 3m; then 21x cycles of 98C for 20s, 61C for 20s, and 72C for 30s; and, finally, 72C for 3m. Reaction products were pooled and gel extracted by loading a 1% TAE gel, excising the 300 bp band, and purifying using a Zymo Research gel extraction kit. Each of the 2 extraction products (N-terminal and C-terminal) were then amplified for 23x cycles each using the same protocol in order to generate sufficient DNA for downstream reactions. To generate concatenations, 5 uL of 15 ng/uL product from each product, N-terminal and C-terminal, were mixed with 4 uL T4 buffer (NEB B0202S), 4 uL BbsI-HF (NEB R3539L, 1 uL T4 ligase (NEB M0202M), and 1uL of an XTEN linker-encoding DNA amplicon. This linker was encoded with variable nucleotides in codon wobble positions to permit variable DNA sequence in between domains and prevent lentiviral recombination without varying the amino acid sequence of the linker. To complete the GoldenGate reaction, the mix was thermocycled for 65x cycles at 37C for 5m, then 16C for 5m, before a series of incubations at 37C for 1h, 65C for 20m, and 16C for 1h. The reaction product was run on a 1% TAE gel and the 600bp band of domain pair concatenations was extracted with a Zymo gel extraction kit. The concatenation product was then PCR amplified as before, using the outer primers, for 15x cycles. Amplification products were then split into 12x GoldenGate reactions, each consisting of 75ng (1.33 uL) Bsmbi-v2 digested pJT039 lentiviral backbone, 10 ng (0.64 uL) concatenation amplicon, 2 uL T4 buffer, 1 uL BsmbI-v2 Golden Gate Assembly Kit (NEB E1602L), and 15.03 uL nuclease free H2O. Reactions were thermocycled with 65x cycles at 42C for 5m and 16C for 5m, before a final incubation at 42C for 5m and 70C for 20m. The reactions were then pooled and purified with a QIAgen MinElute column, eluting in 6 uL ddH2O. 2 uL of eluent was then electroporated into each of 2 tubes of 50 uL Endura electrocompetent cells (Lucigen, Cat#60242-2) as per manufacturer’s instructions. Cells were then plated onto 4x large 10” x 10” LB plates with carbenicillin. After overnight growth colonies were scraped into a collection bottle and plasmid pools were extracted using a plasmid maxiprep kit (Qiagen #12662). 2 smaller plates of LB with carbenicillin were also prepared with 1:20 diluted cells to count colonies and confirm transformation efficiency. Approximately 50 colonies were Sanger sequenced (Quintara) to estimate cloning efficiencies and the proportion of empty backbone plasmids in the pool.

### Co-recruit screen

We plated 15 × 10^6^ HEK293T cells on each of 10x 15-cm tissue culture plates in 30 mL DMEM, grew them overnight, and then transfected each with 8 ug of an equimolar mixture of three third-generation packaging plasmids (pMD2.G, psPAX2, pMDLg/pRRE) and 8 ug of rTetR-concatenation library vectors using 50 mL of polyethylenimine (PEI, Polysciences #23966). Packaging plasmids were gifts from Didier Trono. Lentivirus was harvested after 48 and 72 hours of incubation and filtered through a 0.45 mm PVDF filter (Millipore) to remove debris. K562 cells were infected with lentivirus via spinfection for 2 hours, with 2 replicates each spinfected separately. After 48 hours of growth, infected cells were selected with 10 mg/mL blasticidin (Gibco). Infection and selection efficiency were monitored each day with flow cytometry using a Biorad ZE5 flow cytometer. Infection coverage was approximately 600x for each replicate, while maintenance coverage was maintained between 10,000 and 20,000x cells per library element. On day 8 post-infection, cells were treated with 1000 ng/mL doxycycline (Fisher Scientific) for 2 days for activation or 5 days for repression.

### Co-recruit library prep and sequencing

Genomic DNA was extracted with the QIAgen Blood Maxi Kit following the manufacturer’s instructions with up to 1 × 10^8^ cells per column. Domain sequences were amplified by PCR with primers containing Illumina adapters as extensions. 27-55x 100uL PCRs were set up on ice, with the genomic DNA available for each experiment dictating the number of reactions. 10 ug of genomic DNA, 0.5 uL of each 100M primer, and 50uL of NEBnext Ultra 2x Master Mix (NEB) was used in each reaction. Samples were thermocycled at 98C for 3m; 33 cycles of 98C for 10s, 63C for 1m, 72C for 30s; and 72C for 2m. PCR reactions were pooled and run on a 1% TAE gel, the 600bp band was excised, and DNA was purified using QIAquick gel extraction kit (Qiagen) and eluted into nonstick tubes (Ambion). Samples were sequenced on an Illumina HiSeq (2×150bp, Admera Health).

### Computing co-recruit enrichment scores

Demultiplexed sequencing reads were provided by Admera. Domain 1 and domain 2 reads were aligned separately using bowtie2, trimming 30 base pairs from both 5’ and 3’ ends, and were then each paired read was identified as a particular library member using a python script iterating over each read pair. The total number of reads for each library member was then summed up using another python script. Library members were required to have at least 5 ON reads or 5 OFF reads as well as 50 total reads across both ON and OFF subpopulations to be considered for downstream analysis. For every qualifying library member, the ON or OFF read count was then set to 5 if it was less than 5. Then, ON read counts were normalized by dividing each library member’s read count by the total number of ON reads, and the same was done for OFF counts. Overall enrichment scores were then computed by taking the log_2_ of the ratio of normalized ON read counts to normalized OFF read counts.

### Labeling domains as activators, repressors, and duals

We labeled a domain as an activator if the following 2 conditions held for all concatenations including that domain and a negative control: 1) at least 19% functioned as activators (i.e. for each, activation log_2_(ON:OFF) > −0.0236); 2) the average activation log_2_(ON:OFF) for all pairs was greater than or equal to −0.5, or the activation log_2_(ON:OFF) for all pairs that functioned as activators was greater than or equal to 0.95. We labeled a domain as a repressor if the following 2 conditions held for all concatenations including that domain and a negative control: 1) at least 19% functioned as repressors (i.e. repression log_2_(ON:OFF) < 0.771); 2) the average repression log_2_(ON:OFF) for all pairs was less than or equal to −0.3, or the repression log_2_(ON:OFF) for all pairs that functioned as repressors was less than or equal to 0.20. Domains that functioned as both activators and repressors were labeled as duals. Domains that functioned as neither were labeled as non-hits.

### KRAB-Nuclear Pfam screen cloning and cell culture

An initial backbone was generated by digesting pJT126 (AddGene #161926) with BsmbI, then ligating in a sequence corresponding to the ZNF10 KRAB domain. Then, the nuclear Pfam library was cloned into the downstream GoldenGate cloning site, and used to produce lentivirus identically as in^21^. K562 cells were then spinfected in 2 separate replicates with this KRAB+nuclear Pfam library lentivirus at an average MOI of 0.015, corresponding to an infection coverage of 85x per replicate. 72 hours post-infection 10 mg/mL blasticidin (Gibco) was added. 48 hours after blasticidin addition, 100 ng/mL doxycycline was added and cells were maintained in both blast + dox for 5 more days. Cells were sampled at day 2 of doxycycline recruitment for magnetic separation and downstream sequencing. After 5 days of recruitment, cells were spun down, washed with PBS, and resuspended in dox-free/blast-free RPMI. 5 days after doxycycline release, cells were again sampled for magnetic separation and downstream sequencing. Cells were maintained at a maintenance coverage of 3,000-30,000 per library element on average. Library preparation and sequencing was performed identically as in to sequence the variable effector domain ^21^

### Magnetic Separation

Cells to be separated were spun down at 300 x *g* for 5m and media was aspirated. Cells were then resuspended in PBS (Gibco), spun down again, and PBS was aspirated in order to wash the cells. Dynabeads Protein G were resuspended by vortexing for 30 seconds. 50mL of blocking buffer was prepared per 2 × 10^8^ cells by adding 1 g biotin-free BSA (Sigma Aldrich) and 200 mL of 0.5 M pH 8.0 EDTA into DPBS (GIBCO), vacuum filtering with a 0.22-mm filter (Millipore), and keeping on ice. For all screens 60 uL of beads were used per 10 million cells. Magnetic separation was otherwise performed as in^21^.

### Low-throughput validations

Plasmids for each validation were produced using GoldenGate cloning as described in^21^. Then, 750ng plasmid and 750ng pack mix were mixed in 200uL serum-free OptiMEM (Gibco) along with 2-8uL PEI. Meanwhile, HEK293T cells were plated in 5mL DMEM supplemented with 10% FBS and 1% PSG in one well of a 6-well plate (Fisher Scientific) for each validation. The plasmid-pack mix-optiMEM mixture was added to the HEK293T cells to initiate lentivirus production, and virus was harvested 48 and 72 hours post incubation and filtered through a 0.45mm PDMS filter. For each validation, 2.50 × 10^5^ K562 cells were spinfected with 1mL virus for 2 hours, and 10 mg/mL blasticidin was added after 48 hours. After 7 days of blasticidin selection, cells were spun down and resuspended in RPMI with doxycycline for recruitment.

### Model of transcriptional repression

In our model, λ defines the amount of mRNA in Active (ON) cells. We used the experimental data to estimate these parameters for our synthetic reporter. We directly fit the mean and standard deviation of fluorescence of Silent (OFF) cells from experimental data. We assumed that a population of cells each containing *m* mRNA molecules produced a distribution of log_10_ mCitrine-A fluorescence values centered at (*m*/600) + 6.5, and directly fit the standard deviation of this population from experimental data. We constrained mRNA levels to range between 0 and 1500 molecules per cell. Thus, the probability of seeing a cell produce a log_10_ mCitrine-A fluorescence level of *c* given parameters σ_ON_, μ_OFF_, σ_OFF_, and λ is given by the equations:

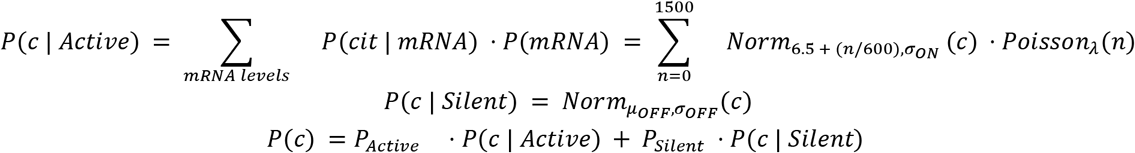

where P_Active_ and P_Silent_ are the probability that a given cell is active or silent, respectively. Parameters were fit directly to data using scipy.optimize.curve_fit. Note that P_Active_ + P_Silent_ = 1 and that when t ≤ t_lag_, P_Silent_ = 0.

To measure silencing, for every day of recruitment, 2000 cells were sampled to ensure equal sampling of log_10_ mCitrine-A fluorescence intensity across recruitment timepoints. The fraction of background silenced cells was directly computed using the rTetR-only recruitment data, with cells being labeled as Silent if their mCitrine-A fluorescence value was less than 10^7^. Then, parameters μ_OFF_ and σ_OFF_ were directly estimated by examining the distribution of cells with log fluorescence values less than 7. λ was determined by estimating the mean amount of mRNA produced in ON cells at day 0, using the above mapping between mRNA counts and fluorescence values. k_s_ and t_lag_ were fit by examining the fraction of cells silenced over time; we assumed that the fraction of Active cells decreased with rate k_s_ after t_lag_ time had elapsed^10,49^. Finally, γ was determined by fitting a 2-state Gaussian Mixture model to fluorescence data for each day and fitting a line to the locations of the higher-fluorescence peak; we assumed λ decays linearly after silencing with slope γ.

### Flow cytometry analysis

Flow cytometry analysis was performed in Python using Cytoflow^78^. For all flow experiments, live cells were gated using Cytoflow’s DensityGateOp, keeping 90% of cells, and then thresholded for mCherry+ cells using the ThresholdGateOp. To determine the fraction of cells that were on, we computed the fraction of cells with log_10_ mCitrine-A fluorescence greater than 7 as measured on a BioRad ZE5 flow cytometer. Normalization of mCitrine levels to the no-dox condition was performed as follows: *f_off,norm_* = (*f_off,dox_* – *f_off,no dox_*) ÷ (1 – *f_off.no dox_*) where f_off_ denotes the fraction of cells off for any given condition. Cytometry analysis was otherwise performed identically as in^20,21^.

## Data and code availability

All raw NGS data and associated processed data generated in this study will be deposited in the NCBI GEO database upon publication. All data besides raw NGS data used for this manuscript, and all code, are available at Zenodo at DOI 10.5281/zenodo.7453682.

## Acknowledgements

We would like to thank Michaela Hinks, Jennifer Gucwa, Nicole DelRosso, Abby Thurm, David Yao, Priyanka Shrestha, Zeppelin Cat, Ophelia Hinks, Shivam Verma, Ben Doughty, Mary Frances Gallagher, Kaushik Ragunathan, Zara Weinberg, and all members of the Bintu lab for helpful conversations and assistance. We would like to thank Anshul Kundaje and Will Greenleaf for valuable feedback throughout the project. AXM was supported by Stanford University Medical Scientist Training Program grants T32-GM007365 and T32-GM145402, and a Stanford Bio-X SIGF. JT is supported by the F99/K00 fellowship of the National Institutes of Health (NIH-1F99DK126120-01; NIH-4K00DK126120-03). SAR is supported by an NSF GRFP (DGE-1656518). MCB is supported by a grant from Stanford ChEM-H and an NIH Director’s New Innovator Award (1DP2HD08406901). This work was supported by BWF CASI (LB), NIH-NIGMS R35M128947 (LB), and NIH-NIGRI R01HG011866 (LB and MCB).

## Author Contributions

AXM and LB designed the study with significant intellectual contributions from JT. AXM and JT designed the combinatorial recruitment library and performed the KRAB-Pfam screen. KS cloned the KRAB-Pfam library. AXM performed the combinatorial recruitment screen with help from CL. AXM, SA, SAR, and CA all performed individual recruitment assay experiments. AXM performed data analysis with input from LB, JT, and MCB. AXM and LB wrote the manuscript, with contributions and feedback from all authors. LB and MCB supervised the project.

## CRediT Statement

AXM: conceptualization, methodology, software, formal analysis, investigation, data curation, writing - original draft, writing - review & editing, visualization. JT: conceptualization, methodology, investigation, writing - review & editing. SA: investigation, writing - review & editing. SR: investigation, writing - review & editing. CA: investigation, writing - review & editing. CL: investigation, writing - review & editing. MCB: conceptualization, resources, writing - review & editing, supervision, project administration. LB: conceptualization, methodology, resources, writing - review & editing, supervision, project administration, funding acquisition.

## Additional Information

Supplementary information is available for this paper. Correspondence and requests for materials should be addressed to.

## References

1. Frietze, S., and Farnham, P.J. (2011). Transcription Factor Effector Domains. In A Handbook of Transcription Factors, T. R. Hughes, ed. (Springer Netherlands), pp. 261–277. 10.1007/978-90-481-9069-0_12.

2. Soto, L.F., Li, Z., Santoso, C.S., Berenson, A., Ho, I., Shen, V.X., Yuan, S., and Fuxman Bass, J.I. (2021). Compendium of human transcription factor effector domains. Mol. Cell. 10.1016/j.molcel.2021.11.007.

3. Lambert, S.A., Jolma, A., Campitelli, L.F., Das, P.K., Yin, Y., Albu, M., Chen, X., Taipale, J., Hughes, T.R., and Weirauch, M.T. (2018). The Human Transcription Factors. Cell 172, 650–665. 10.1016/j.cell.2018.01.029.

4. Stegmaier, P., Kel, A.E., and Wingender, E. (2004). Systematic DNA-binding domain classification of transcription factors. Genome Inform. 15, 276–286.

5. Harrison, S.C. (1991). A structural taxonomy of DNA-binding domains. Nature 353, 715–719. 10.1038/353715a0.

6. Hathaway, N.A., Bell, O., Hodges, C., Miller, E.L., Neel, D.S., and Crabtree, G.R. (2012). Dynamics and memory of heterochromatin in living cells. Cell 149, 1447–1460. 10.1016/j.cell.2012.03.052.

7. Keung, A.J., Bashor, C.J., Kiriakov, S., Collins, J.J., and Khalil, A.S. (2014). Using targeted chromatin regulators to engineer combinatorial and spatial transcriptional regulation. Cell 158, 110–120. 10.1016/j.cell.2014.04.047.

8. Kungulovski, G., Nunna, S., Thomas, M., Zanger, U.M., Reinhardt, R., and Jeltsch, A. (2015). Targeted epigenome editing of an endogenous locus with chromatin modifiers is not stably maintained. Epigenetics Chromatin 8, 12. 10.1186/s13072-015-0002-z.

9. Amabile, A., Migliara, A., Capasso, P., Biffi, M., Cittaro, D., Naldini, L., and Lombardo, A. (2016). Inheritable Silencing of Endogenous Genes by Hit-and-Run Targeted Epigenetic Editing. Cell 167, 219–232.e14. 10.1016/j.cell.2016.09.006.

10. Bintu, L., Yong, J., Antebi, Y.E., McCue, K., Kazuki, Y., Uno, N., Oshimura, M., and Elowitz, M.B. (2016). Dynamics of epigenetic regulation at the single cell level. Science 351, 720–724.

11. Tycko, J., Van, M.V., Elowitz, M.B., and Bintu, L. (2017). Advancing towards a global mammalian gene regulation model through single-cell analysis and synthetic biology. Current Opinion in Biomedical Engineering 4, 174–193. 10.1016/j.cobme.2017.10.011.

12. O’Geen, H., Ren, C., Nicolet, C.M., Perez, A.A., Halmai, J., Le, V.M., MacKay, J.P., Farnham, P.J., and Segal, D.J. (2017). DCas9-based epigenome editing suggests acquisition of histone methylation is not sufficient for target gene repression. Nucleic Acids Res. 45, 9901–9916. 10.1093/nar/gkx578.

13. Sanborn, A.L., Yeh, B.T., Feigerle, J.T., Hao, C.V., Townshend, R.J., Lieberman Aiden, E., Dror, R.O., and Kornberg, R.D. (2021). Simple biochemical features underlie transcriptional activation domain diversity and dynamic, fuzzy binding to Mediator. Elife 10. 10.7554/eLife.68068.

14. Lee, J.B., Caywood, L.M., Lo, J.Y., Levering, N., and Keung, A.J. (2021). Mapping the dynamic transfer functions of eukaryotic gene regulation. Cell Syst 12, 1079–1093.e6. 10.1016/j.cels.2021.08.003.

15. Chasman, D.I., Leatherwood, J., Carey, M., Ptashne, M., and Kornberg, R.D. (1989). Activation of yeast polymerase II transcription by herpesvirus VP16 and GAL4 derivatives in vitro. Mol. Cell. Biol. 9, 4746–4749. 10.1128/mcb.9.11.4746-4749.1989.

16. Furth, N., and Shema, E. (2022). It’s all in the combination: decoding the epigenome for cancer research and diagnostics. Curr. Opin. Genet. Dev. 73, 101899. 10.1016/j.gde.2022.101899.

17. Wolberger, C. (1998). Combinatorial transcription factors. Curr. Opin. Genet. Dev. 8, 552–559. 10.1016/s0959-437x(98)80010-5.

18. Ravasi, T., Suzuki, H., Cannistraci, C.V., Katayama, S., Bajic, V.B., Tan, K., Akalin, A., Schmeier, S., Kanamori-Katayama, M., Bertin, N., et al. (2010). An atlas of combinatorial transcriptional regulation in mouse and man. Cell 140, 744–752. 10.1016/j.cell.2010.01.044.

19. Reiter, F., Wienerroither, S., and Stark, A. (2017). Combinatorial function of transcription factors and cofactors. Curr. Opin. Genet. Dev. 43, 73–81. 10.1016/j.gde.2016.12.007.

20. DelRosso, N., Tycko, J., Suzuki, P., Andrews, C., Aradhana, Mukund, A., Liongson, I., Ludwig, C., Spees, K., Fordyce, P., et al. (2022). Large-scale mapping and systematic mutagenesis of human transcriptional effector domains. bioRxiv, 2022.08.26.505496. 10.1101/2022.08.26.505496.

21. Tycko, J., DelRosso, N., Hess, G.T., Aradhana, Banerjee, A., Mukund, A., Van, M.V., Ego, B.K., Yao, D., Spees, K., et al. (2020). High-Throughput Discovery and Characterization of Human Transcriptional Effectors. Cell 183, 2020–2035.e16. 10.1016/j.cell.2020.11.024.

22. Losson, R., and Nielsen, A.L. (2010). The NIZP1 KRAB and C2HR domains cross-talk for transcriptional regulation. Biochimica et Biophysica Acta - Gene Regulatory Mechanisms 1799, 463–468. 10.1016/j.bbagrm.2010.02.003.

23. Beerli, R.R., Segal, D.J., Dreier, B., and Barbas, C.F., 3rd (1998). Toward controlling gene expression at will: specific regulation of the erbB-2/HER-2 promoter by using polydactyl zinc finger proteins constructed from modular building blocks. Proc. Natl. Acad. Sci. U. S. A. 95, 14628–14633. 10.1073/pnas.95.25.14628.

24. Chavez, A., Scheiman, J., Vora, S., Pruitt, B.W., Tuttle, M., P R Iyer, E., Lin, S., Kiani, S., Guzman, C.D., Wiegand, D.J., et al. (2015). Highly efficient Cas9-mediated transcriptional programming. Nat. Methods 12, 326–328. 10.1038/nmeth.3312.

25. Nuñez, J.K., Chen, J., Pommier, G.C., Cogan, J.Z., Replogle, J.M., Adriaens, C., Ramadoss, G.N., Shi, Q., Hung, K.L., Samelson, A.J., et al. (2021). Genome-wide programmable transcriptional memory by CRISPR-based epigenome editing. Cell 184, 2503–2519.e17. 10.1016/j.cell.2021.03.025.

26. Thakore, P.I., Black, J.B., Hilton, I.B., and Gersbach, C.A. (2016). Editing the epigenome: technologies for programmable transcription and epigenetic modulation. Nature Methods. 10.1038/nmeth.3733.

27. Holtzman, L., and Gersbach, C.A. (2018). Editing the Epigenome: Reshaping the Genomic Landscape. Annu. Rev. Genomics Hum. Genet. 19, 43–71. 10.1146/annurev-genom-083117-021632.

28. Black, J.B., and Gersbach, C.A. (2018). Synthetic transcription factors for cell fate reprogramming. Curr. Opin. Genet. Dev. 52, 13–21. 10.1016/j.gde.2018.05.001.

29. Elkon, R., and Agami, R. (2017). Characterization of noncoding regulatory DNA in the human genome. Nat. Biotechnol. 35, 732–746. 10.1038/nbt.3863.

30. Erijman, A., Kozlowski, L., Sohrabi-Jahromi, S., Fishburn, J., Warfield, L., Schreiber, J., Noble, W.S., Söding, J., and Hahn, S. (2020). A High-Throughput Screen for Transcription Activation Domains Reveals Their Sequence Features and Permits Prediction by Deep Learning. Molecular Cell 79, 1066. 10.1016/j.molcel.2020.08.013.

31. Staller, M.V., Holehouse, A.S., Swain-Lenz, D., Das, R.K., Pappu, R.V., and Cohen, B.A. (2018). A High-Throughput Mutational Scan of an Intrinsically Disordered Acidic Transcriptional Activation Domain. Cell Syst 6, 444–455.e6. 10.1016/j.cels.2018.01.015.

32. Arnold, C.D., Nemčko, F., Woodfin, A.R., Wienerroither, S., Vlasova, A., Schleiffer, A., Pagani, M., Rath, M., and Stark, A. (2018). A high-throughput method to identify trans-activation domains within transcription factor sequences. EMBO J. 37, e98896. 10.15252/embj.201798896.

33. Alerasool, N., Leng, H., Lin, Z.-Y., Gingras, A.-C., and Taipale, M. (2022). Identification and functional characterization of transcriptional activators in human cells. Mol. Cell 82, 677–695.e7. 10.1016/j.molcel.2021.12.008.

34. Staller, M.V., Ramirez, E., Kotha, S.R., Holehouse, A.S., Pappu, R.V., and Cohen, B.A. (2022). Directed mutational scanning reveals a balance between acidic and hydrophobic residues in strong human activation domains. Cell Syst 13, 334–345.e5. 10.1016/j.cels.2022.01.002.

35. Gao, Y., Xiong, X., Wong, S., Charles, E.J., Lim, W.A., and Qi, L.S. (2016). Complex transcriptional modulation with orthogonal and inducible dCas9 regulators. Nat. Methods 13, 1043–1049. 10.1038/nmeth.4042.

36. Cura, V., and Cavarelli, J. (2021). Structure, Activity and Function of the PRMT2 Protein Arginine Methyltransferase. Life 11. 10.3390/life11111263.

37. Pak, M.L., Lakowski, T.M., Thomas, D., Vhuiyan, M.I., Hüsecken, K., and Frankel, A. (2011). A protein arginine N-methyltransferase 1 (PRMT1) and 2 heteromeric interaction increases PRMT1 enzymatic activity. Biochemistry 50, 8226–8240. 10.1021/bi200644c.

38. Hou, W., Nemitz, S., Schopper, S., Nielsen, M.L., Kessels, M.M., and Qualmann, B. (2018). Arginine Methylation by PRMT2 Controls the Functions of the Actin Nucleator Cobl. Dev. Cell 45, 262–275.e8. 10.1016/j.devcel.2018.03.007.

39. Vhuiyan, M.I., Pak, M.L., Park, M.A., Thomas, D., Lakowski, T.M., Chalfant, C.E., and Frankel, A. (2017). PRMT2 interacts with splicing factors and regulates the alternative splicing of BCL-X. J. Biochem. 162, 17–25. 10.1093/jb/mvw102.

40. Kremerskothen, J., Plaas, C., Büther, K., Finger, I., Veltel, S., Matanis, T., Liedtke, T., and Barnekow, A. (2003). Characterization of KIBRA, a novel WW domain-containing protein. Biochem. Biophys. Res. Commun. 300, 862–867. 10.1016/s0006-291x(02)02945-5.

41. Xiao, L., Chen, Y., Ji, M., and Dong, J. (2011). KIBRA regulates Hippo signaling activity via interactions with large tumor suppressor kinases. J. Biol. Chem. 286, 7788–7796. 10.1074/jbc.M110.173468.

42. Gordon, W.R., Vardar-Ulu, D., Histen, G., Sanchez-Irizarry, C., Aster, J.C., and Blacklow, S.C. (2007). Structural basis for autoinhibition of Notch. Nat. Struct. Mol. Biol. 14, 295–300. 10.1038/nsmb1227.

43. Bray, S.J. (2016). Notch signalling in context. Nat. Rev. Mol. Cell Biol. 17, 722–735. 10.1038/nrm.2016.94.

44. Raj, A., Peskin, C.S., Tranchina, D., Vargas, D.Y., and Tyagi, S. (2006). Stochastic mRNA synthesis in mammalian cells. PLoS Biol. 4, e309. 10.1371/journal.pbio.0040309.

45. Friedman, N., Cai, L., and Xie, X.S. (2006). Linking stochastic dynamics to population distribution: an analytical framework of gene expression. Phys. Rev. Lett. 97, 168302. 10.1103/PhysRevLett.97.168302.

46. Dar, R.D., Razooky, B.S., Singh, A., Trimeloni, T.V., McCollum, J.M., Cox, C.D., Simpson, M.L., and Weinberger, L.S. (2012). Transcriptional burst frequency and burst size are equally modulated across the human genome. Proc. Natl. Acad. Sci. U. S. A. 109, 17454–17459. 10.1073/pnas.1213530109.

47. Peccoud, J., and Ycart, B. (1995). Markovian modeling of gene-product synthesis. Theoretical Population Biology 48, 222–234. 10.1006/tpbi.1995.1027.

48. Hao, N., and O’Shea, E.K. (2012). Signal-dependent dynamics of transcription factor translocation controls gene expression. Nat. Struct. Mol. Biol. 19, 31–40. 10.1038/nsmb.2192.

49. Mukund, A., and Bintu, L. (2022). Temporal signaling, population control, and information processing through chromatin-mediated gene regulation. J. Theor. Biol. 535, 110977. 10.1016/j.jtbi.2021.110977.

50. Martinez-Corral, R., Park, M., Biette, K., Friedrich, D., Scholes, C., Khalil, A.S., Gunawardena, J., and DePace, A.H. (2021). Transcriptional kinetic synergy: a complex landscape revealed by integrating modelling and synthetic biology. bioRxiv, 2020.08.31.276261. 10.1101/2020.08.31.276261.

51. Park, H., Wahl, M.I., Afar, D.E., Turck, C.W., Rawlings, D.J., Tam, C., Scharenberg, A.M., Kinet, J.P., and Witte, O.N. (1996). Regulation of Btk function by a major autophosphorylation site within the SH3 domain. Immunity 4, 515–525. 10.1016/s1074-7613(00)80417-3.

52. Mohamed, A.J., Vargas, L., Nore, B.F., Backesjo, C.M., Christensson, B., and Smith, C.I. (2000). Nucleocytoplasmic shuttling of Bruton’s tyrosine kinase. J. Biol. Chem. 275, 40614–40619. 10.1074/jbc.M006952200.

53. Lee, Y.-T., Ayoub, A., Park, S.-H., Sha, L., Xu, J., Mao, F., Zheng, W., Zhang, Y., Cho, U.-S., and Dou, Y. (2021). Mechanism for DPY30 and ASH2L intrinsically disordered regions to modulate the MLL/SET1 activity on chromatin. Nat. Commun. 12, 2953. 10.1038/s41467-021-23268-9.

54. Tando, T., Ishizaka, A., Watanabe, H., Ito, T., Iida, S., Haraguchi, T., Mizutani, T., Izumi, T., Isobe, T., Akiyama, T., et al. (2010). Requiem protein links RelB/p52 and the Brm-type SWI/SNF complex in a noncanonical NF-κB pathway. J. Biol. Chem. 285, 21951–21960.

55. Kim, G.-D., Ni, J., Kelesoglu, N., Roberts, R.J., and Pradhan, S. (2002). Co-operation and communication between the human maintenance and de novo DNA (cytosine-5) methyltransferases. EMBO J. 21, 4183–4195. 10.1093/emboj/cdf401.

56. Larson, M.H., Gilbert, L.A., Wang, X., Lim, W.A., Weissman, J.S., and Qi, L.S. (2013). CRISPR interference (CRISPRi) for sequence-specific control of gene expression. Nat. Protoc. 8, 2180–2196. 10.1038/nprot.2013.132.

57. Jacobs, C.L., Badiee, R.K., and Lin, M.Z. (2018). StaPLs: versatile genetically encoded modules for engineering drug-inducible proteins. Nat. Methods 15, 523–526. 10.1038/s41592-018-0041-z.

58. Reményi, A., Schöler, H.R., and Wilmanns, M. (2004). Combinatorial control of gene expression. Nat. Struct. Mol. Biol. 11, 812–815. 10.1038/nsmb820.

59. Zeitlinger, J. (2020). Seven myths of how transcription factors read the cis-regulatory code. Curr Opin Syst Biol 23, 22–31. 10.1016/j.coisb.2020.08.002.

60. Charest, J., Daniele, T., Wang, J., Bykov, A., Mandlbauer, A., Asparuhova, M., Röhsner, J., Gutiérrez-Pérez, P., and Cochella, L. (2020). Combinatorial Action of Temporally Segregated Transcription Factors. Dev. Cell 55, 483–499.e7. 10.1016/j.devcel.2020.09.002.

61. Bashor, C.J., Patel, N., Choubey, S., Beyzavi, A., Kondev, J., Collins, J.J., and Khalil, A.S. (2019). Complex signal processing in synthetic gene circuits using cooperative regulatory assemblies. Science 364, 593–597. 10.1126/science.aau8287.

62. Roybal, K.T., Rupp, L.J., Morsut, L., Walker, W.J., McNally, K.A., Park, J.S., and Lim, W.A. (2016). Precision Tumor Recognition by T Cells With Combinatorial Antigen-Sensing Circuits. Cell 164, 770–779. 10.1016/j.cell.2016.01.011.

63. Weinberg, Z.Y., Hilburger, C.E., Kim, M., Cao, L., Khalid, M., Elmes, S., Diwanji, D., Hernandez, E., Lopez, J., Schaefer, K., et al. (2021). Sentinel cells enable genetic detection of SARS-CoV-2 Spike protein. bioRxiv. 10.1101/2021.04.20.440678.

64. Zhu, I., Liu, R., Garcia, J.M., Hyrenius-Wittsten, A., Piraner, D.I., Alavi, J., Israni, D.V., Liu, B., Khalil, A.S., and Roybal, K.T. (2022). Modular design of synthetic receptors for programmed gene regulation in cell therapies. Cell 185, 1431–1443.e16. 10.1016/j.cell.2022.03.023.

65. Lee, S., Khalil, A.S., and Wong, W.W. (2022). Recent progress of gene circuit designs in immune cell therapies. Cell Syst 13, 864–873. 10.1016/j.cels.2022.09.006.

66. Konermann, S., Brigham, M.D., Trevino, A.E., Joung, J., Abudayyeh, O.O., Barcena, C., Hsu, P.D., Habib, N., Gootenberg, J.S., Nishimasu, H., et al. (2015). Genome-scale transcriptional activation by an engineered CRISPR-Cas9 complex. Nature 517, 583–588. 10.1038/nature14136.

67. Schüle, R., Rangarajan, P., Kliewer, S., Ransone, L.J., Bolado, J., Yang, N., Verma, I.M., and Evans, R.M. (1990). Functional antagonism between oncoprotein c-Jun and the glucocorticoid receptor. Cell 62, 1217–1226. 10.1016/0092-8674(90)90397-w.

68. Gill, G., and Ptashne, M. (1988). Negative effect of the transcriptional activator GAL4. Nature 334, 721–724. 10.1038/334721a0.

69. Al-Radhawi, M.A., Ali Al-Radhawi, M., Del Vecchio, D., and Sontag, E.D. (2019). Multi-modality in gene regulatory networks with slow promoter kinetics. PLOS Computational Biology 15, e1006784. 10.1371/journal.pcbi.1006784.

70. Mamrak, N.E., Alerasool, N., Griffith, D., Holehouse, A.S., Taipale, M., and Lionnet, T. (2022). The kinetic landscape of human transcription factors. bioRxiv, 2022.06.01.494187. 10.1101/2022.06.01.494187.

71. Lim, W.A. (2022). The emerging era of cell engineering: Harnessing the modularity of cells to program complex biological function. Science 378, 848–852. 10.1126/science.add9665.

72. Majello, B., De Luca, P., and Lania, L. (1997). Sp3 is a bifunctional transcription regulator with modular independent activation and repression domains. J. Biol. Chem. 272, 4021–4026. 10.1074/jbc.272.7.4021.

73. Jacobs, J., Pagani, M., Wenzl, C., and Stark, A. (2022). Widespread regulatory specificities between transcriptional corepressors and enhancers in Drosophila. bioRxiv, 2022.11.07.515017. 10.1101/2022.11.07.515017.

74. Shakiba, N., Jones, R.D., Weiss, R., and Del Vecchio, D. (2021). Context-aware synthetic biology by controller design: Engineering the mammalian cell. Cell Syst 12, 561–592. 10.1016/j.cels.2021.05.011.

75. Dods, G., Gómez-Schiavon, M., El-Samad, H., and Ng, A.H. (2020). Accurate prediction of genetic circuit behavior requires multidimensional characterization of parts. bioRxiv, 2020.05.30.122077. 10.1101/2020.05.30.122077.

76. Baetica, A.-A., Westbrook, A., and El-Samad, H. (2019). Control theoretical concepts for synthetic and systems biology. Current Opinion in Systems Biology 14, 50–57. 10.1016/j.coisb.2019.02.010.

77. Zulkower, V., and Rosser, S. (2020). DNA Chisel, a versatile sequence optimizer. Bioinformatics 36, 4508–4509. 10.1093/bioinformatics/btaa558.

78. Teague, B. (2022). Cytoflow: A Python Toolbox for Flow Cytometry. bioRxiv, 2022.07.22.501078. 10.1101/2022.07.22.501078.

